# An integrated platform for high-throughput phenospace learning of 3D multilineage organoid systems

**DOI:** 10.1101/2025.11.27.690952

**Authors:** Ryo Okuda, Christoph Harmel, Quan Xu, Héloise Mary, Philip Schulz, Linda Steinacher, Elisa D’Arcangelo, Bruno Gjeta, Marina Signer, Irineja Cubela, Marc Bickle, Matthias P. Lutolf, Lauriane Cabon, Ilya Lukonin, J. Gray Camp

## Abstract

Complex multilineage organoid systems lack quantitative phenotyping methods preserving spatial architecture at high throughput. Current approaches compromise biological complexity, spatial resolution, or robust homogeneous multilineage assembly. We establish an integrated experimental-computational platform for high-throughput spatial phenotyping of multilineage organoids through developing a modular tumoroid culture system incorporating pancreatic ductal adenocarcinoma (PDAC) cells and cancer-associated fibroblasts (CAFs) in 384-well format with multiplexed whole-mount imaging. We developed Phenocoder, a machine learning framework combining conditional variational autoencoders with spatial graph analysis to extract multiscale organoid features. Rigorous validation demonstrates robust performance in PDAC tumoroids. The platform identifies pathway modulators that disrupt the fibrotic microenvironment and discovers stroma-dependent vulnerabilities, undetectable in monocultures. Extending to immuno-competent tumoroids, we assess fibrosis modulators in combination with T cell bispecific antibodies, identifying treatments that enhance immune cell proliferation and infiltration inducing cancer cell death, validated in patient-derived explants. This platform establishes a generalizable framework for multilineage organoid phenotyping.

## Introduction

Multilineage organoid systems incorporating epithelial, stromal, immune, vascular, and other cell types hold particular promise for modeling disease microenvironments and capturing context-dependent therapeutic responses (Recaldin et al.; Sharpe et al.; Liu et al., a). However, quantitative phenotyping methods that preserve spatial architecture at high throughput remain limited, as current experimental approaches compromise either biological complexity, spatial resolution, or robust multilineage assembly. Overcoming this gap requires integrated experimental-computational platforms capable of systematically phenotyping complex, multilineage organoid systems at scale while maintaining spatial and compositional resolution.

Pancreatic ductal adenocarcinoma (PDAC) persists as a lethal malignancy, characterized by late diagnosis, aggressive progression, and resistance to current therapies (Dreyer et al.; Siegel et al.). A hallmark of PDAC is a dense, desmoplastic stroma, dominated by cancer-associated fibroblasts (CAFs) and extracellular matrix components that create a complex and tumor-promoting microenvironment (Halbrook et al.). This stromal barrier fosters tumor growth, provides a niche for metastatic potential, and impedes drug delivery and immune cell infiltration, contributing to poor patient outcomes (Monteran et al.; Grünwald et al.; Provenzano et al.; Punovuori et al.). Single-cell transcriptome studies of primary PDAC lesions revealed distinct cancer cell states with varying degrees of epithelial-mesenchymal plasticity, metabolic adaptation, lineage de-differentiation and other malignant states (Hwang et al.; Pei et al.; Chen et al.). In addition, single-cell transcriptomics has illuminated CAF heterogeneity, distinguishing myofibroblastic, inflammatory, and antigen-presenting CAF subtypes with distinct functional properties and prognostic implications (Cheng et al., b; Liu et al., b; Gao et al.). Additional immune, vascular, and neuronal cell types and states co-occur within or surround cancer microenvironments. Integration of PDAC cell atlases have identified prevalent cell states that exist in the majority of PDAC patients (Xu et al., a). These high-resolution insights can inform the development of in vitro models that better recapitulate the cellular diversity observed in patient tumors and provide opportunities for therapeutic targeting.

Cancer cells can be isolated from PDAC primary and metastatic tumor biopsies, and cultivated in vitro as three-dimensional cancer organoids (Boj et al.; Raghavan et al.; Driehuis et al., a; Li et al., b). These primarily cystic models provide patient-specific representations of cancer genetic and epigenetic states, and can be used for high-throughput screening and patient-specific therapy assessment (Zapatero et al.; Driehuis et al., b; Cutrona et al.; Hirt et al.). Most cancer organoid models represent only the epithelial cancer compartment (Verstegen et al.). CAFs can also be simultaneously isolated from the same tumor biopsies, propagated in vitro, and used to reconstitute autologous multilineage, stroma-rich, co-culture tumoroids (Xu et al., a; Langer et al.; Seino et al.; Kim et al., a; Sharpe et al.; Barbazan et al.). Recent benchmarking has demonstrated that tumoroids can recapitulate important cell state features of PDAC primary counterparts that develop over time in co-culture (Xu et al., a). However, due to inherent heterogeneity and phenotypic complexity, it remains a major challenge to scale multilineage tumoroid models for screening therapeutic agents.

Systematic mapping of cellular and tissue phenotypes under different perturbation conditions provides a framework for characterizing phenotypic spaces (phenospaces), enabling quantitative assessment of biological responses across multidimensional feature landscapes (Fleck et al.; Rood et al.). Recent advances in high-content screening (HCS) have extended phenotypic profiling to three-dimensional organoid systems (Lukonin et al.; Beghin et al.). However, the inherent complexity of 3D multicellular structures poses significant analytical challenges. Computational frameworks for spatial cellular neighborhood analysis and approaches for generating digitalized organoid representations have advanced morphological quantification capabilities (Ong et al.; Palla et al.; Albanese et al.; Wahle et al.). Phenotypic organoid embeddings have proven valuable for quantifying treatment effects and mapping developmental trajectories (Metzger et al.; Lukonin et al.), yet obstacles remain to develop unsupervised analysis frameworks that can characterize 3D organoid models and extract biologically meaningful information about complex microenvironments from high-content imaging data.

Here, we integrate automated cell culture, multiplexed staining, tissue clearing, and high-content 3D imaging to deeply phenotype tumoroid growth, composition, and cellular arrangements. We develop Phenocoder, a machine learning framework that combines conditional variational autoencoders with spatial graph analysis to generate latent phenotypic embeddings of tumoroid co-cultures, enabling quantitative assessment of spatial cellular organization dynamics and drug responses. High-throughput perturbation screening identified previously unappreciated druggable targets in the PDAC microenvironment that strongly disrupt cancer-CAF interactions. We incorporate tumor infiltrating lymphocytes (TILs) or activated T cells from peripheral blood mononuclear cells (PBMCs) into tumoroids and show that immunocompetent models can be used to assess combination fibrosis modulation with cancer-targeting T cell bispecific antibodies. This integrated approach to learning multivariate feature tumoroid profiles represents a generalizable framework applicable to other complex multicellular systems.

## Results

### Tumoroid co-culture system for high-throughput phenotyping

We established a high-throughput platform to model cancer-CAF interactions and phenotype PDAC tumoroids (Fig. 1a). Patient-derived (KRAS^+^) cancer organoids were co-cultured for 14 days with matched CAFs from the same individual to generate autologous tumoroids (Table S1). Tumoroids were initialized in 384 micro-well plates and cultured using a customized robotic liquid handling workflow (Extended Data Fig. 1a). Robotic deposition of cancer-CAF-ECM mixtures at central positions within each well enabled consistent initializing conditions, which resulted in reproducible morphogenetic growth over a 14 day period which can be monitored by live imaging (Fig. 1b). From this uniform baseline, the co-cultured cells underwent progressive self-organization, with cancer and CAFs orchestrating the emergence of increasingly complex tissue architectures over time. By day 14, the tumoroids exhibited highly organized morphogenetic structures, with cancer cells expressing Arginase2 (ARG2) surrounded by a Vimentin-positive fibrotic network, reminiscent of primary PDAC tissue (Fig. 1c).

**Figure 1.**
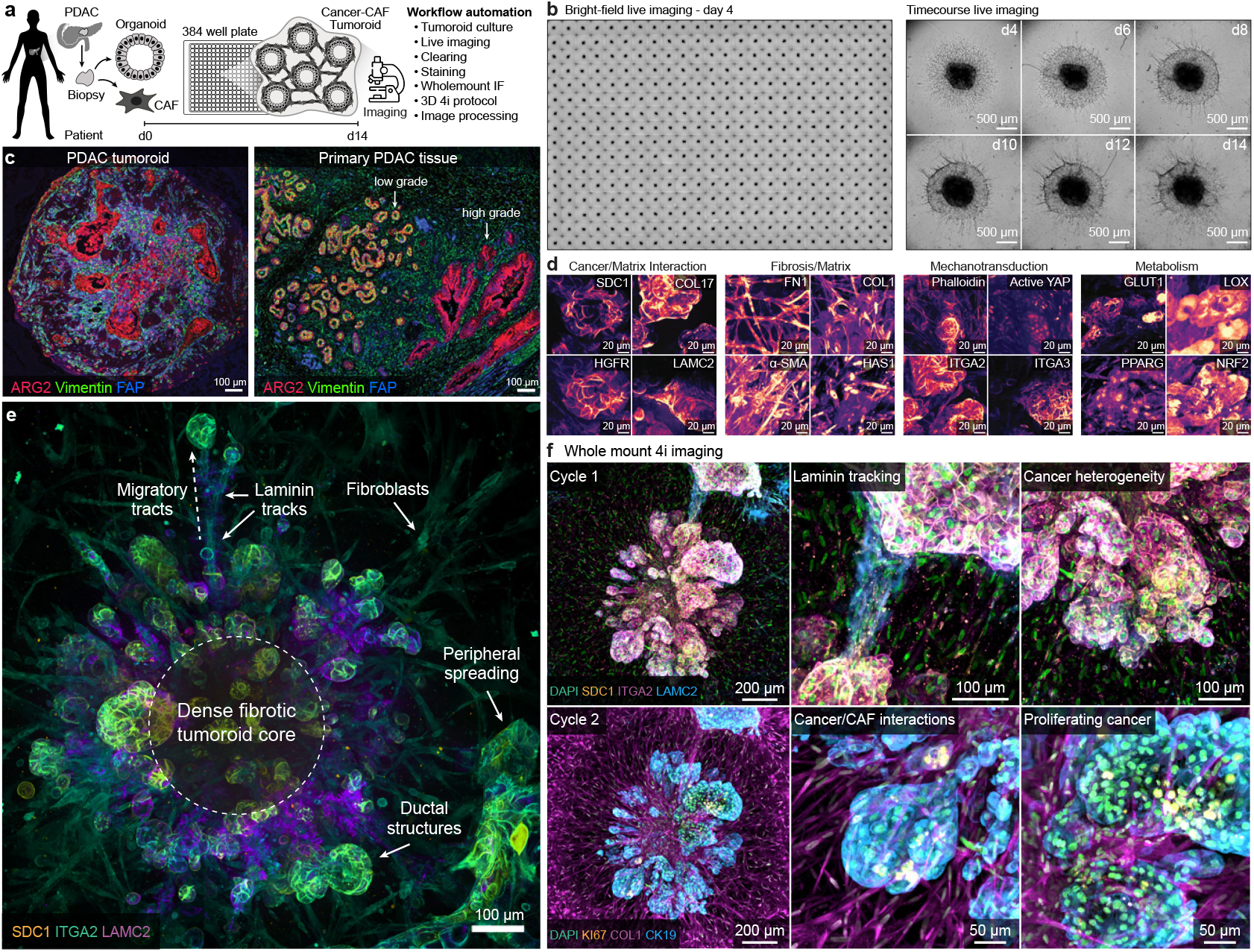
High-throughput phenotyping of 3D human pancreatic tumoroids. a) Overview of the high-content imaging-based phenotypic screening platform for patient-derived PDAC cancer-CAF tumoroids co-cultures. b) Bright field images from day 4 organoids in a 384 well plate (left) and representative tumoroid from selected time points (day 4–14, right). c) Histological analysis of cross-section staining of tumoroids and primary PDAC tissue stained for ARG2, Vimentin, and FAP, highlighting cancer and fibroblastic domains. d) Representative images from high-throughput antibody testing in 384 well plates using automated clearing, staining, and imaging regimes. e) Structural zones and features of the tumoroid stained with SDC1 (orange), ITGA2 (green), and LAMC2 (magenta), revealing distinct architectural regions including migratory tracks, dense fibrotic core, ductal structures, and peripheral spreading. f) Cleared whole-mount 4i imaging of fibrotic tumoroids across two imaging cycles. Cycle 1 shows staining for DAPI (white), SDC1 (orange), ITGA2 (green), and LAMC2 (magenta). Cycle 2 shows DAPI (white), Ki67 (green), COL1 (orange), and CK19 (magenta). Panels highlight laminin tracking, structural heterogeneity, cancer-CAF interactions, and proliferating epithelial regions.

We implemented semi-automated, whole-mount cleared immunofluorescence imaging to screen and assess an array of 35 protein expression patterns, providing an overview of the emergent tissue composition and microenvironment architecture (Fig. 1d, Extended Data Fig. 2a, Table S2). Maximum projection of stacked image acquisition at 20x resolution in cleared whole-mount preparations identified antibody sets that provide information on the ductal structures, extracellular matrix, intracellular signalling status, organelles, and other phenotypic components. Syndecan-1 (SDC1), Phalloidin, and activated YAP revealed progressive architectural remodeling, characterized by dense central cores, structured actin networks, and peripheral zones enriched for YAP signaling-hallmarks of mechanoresponsive tissue organization (Extended Data Fig. 2b). Tumoroids developed spatially organized zones marked by Laminin-*γ*2 (LAMC2), SDC1, and Integrin *α*2 (ITGA2), which localized to ductal epithelium, fibrotic cores, and invasive edges (Fig. 1e). These markers revealed coordinated ECM remodeling and suggested collective cancer cell migration as the tissue matured. Individual z-slices captured laminin-rich tracks and peripheral epithelial morphologies associated with fibroblast networks (Extended Data Fig. 2c). To further characterize the cancer-CAFs interface, we performed cleared 3D imaging of SDC1 with fibronectin (Extended Data Fig. 2d). These data revealed that SDC1^+^ ductal structured cancer cells are not just adjacent to CAF-derived ECM, but are physically enveloped within dense fibronectin networks throughout the tumoroid. 3D-visualization confirmed a spatially resolved architecture, with cancer cells embedded in and overlaid by fibronectin fibrils. High resolution z-slices further demonstrated SDC1 localization at the cancer cell membrane in direct juxtaposition with fibronectin fibers, underscoring the architectural coupling of cancer cells and CAF derived ECM. This organization highlights how CAF derived matrix forms a structural scaffold that integrates and constrains cancer cells, recapitulating hallmarks of PDAC desmoplasia.

Adaptation of the iterative indirect immunofluorescence imaging (4i) method (Gut et al.) to the whole-mount tumoroid imaging setup enabled parallel imaging of tissue architecture markers (SDC1, ITGA2, LAMC2) and proliferation or stromal markers (COL1, Ki67, CK19) within the same specimen through sequential antibody elution and re-staining cycles (Fig. 1f, Extended Data Fig. 1c,d). This approach provided detailed spatial mappings of both structural organization and cellular activity states across the entire 3D tumoroid architecture, with maximum z-projections revealing compartmentalization of proliferative epithelial cells within matrix-rich environments consistent with fibrotic tumor architecture. Altogether, this semi-automated system supports scalable production, culture, live imaging, and endpoint multiplexed whole-mount imaging of tumoroid tissue bearing hallmarks of the primary tissue architecture.

### Reconstruction of spatiotemporal phenotypic dynamics with Phenocoder

We performed multiplexed time-course imaging of 12 antibody sets (Table S2, Extended Data Fig. 3) to register tumoroids (n=480) across 24 markers and DAPI staining at 3, 5, 7, 10, and 14 days of culture, revealing progressive growth over this timeframe (Fig. 2a,b, Extended Data Fig. 4a). The imaging captured temporal transitions towards structured tissue organization, with formation of spatially organized tumor ductal zones, fibrotic stroma, dense tumoroid cores, and migratory fronts, as indicated by SDC1, Phalloidin, and activated YAP expression patterns (Fig. 2b, Extended Data Fig. 3). To quantitatively characterize tumoroid spatiotemporal phenotypic dynamics in an unsupervised manner, we developed Phenocoder (Fig. 2c), a machine learning framework that combines conditional variational autoencoders (cVAEs) with spatial graph analysis to generate latent phenotypic embeddings of complex tissue architectures. Phenocoder integrates both cellular and microenvironmental features across multiple spatial scales, enabling unsupervised, robust, and quantitative assessment of microenvironmental tissue organization in tumoroid co-cultures.

**Figure 2.**
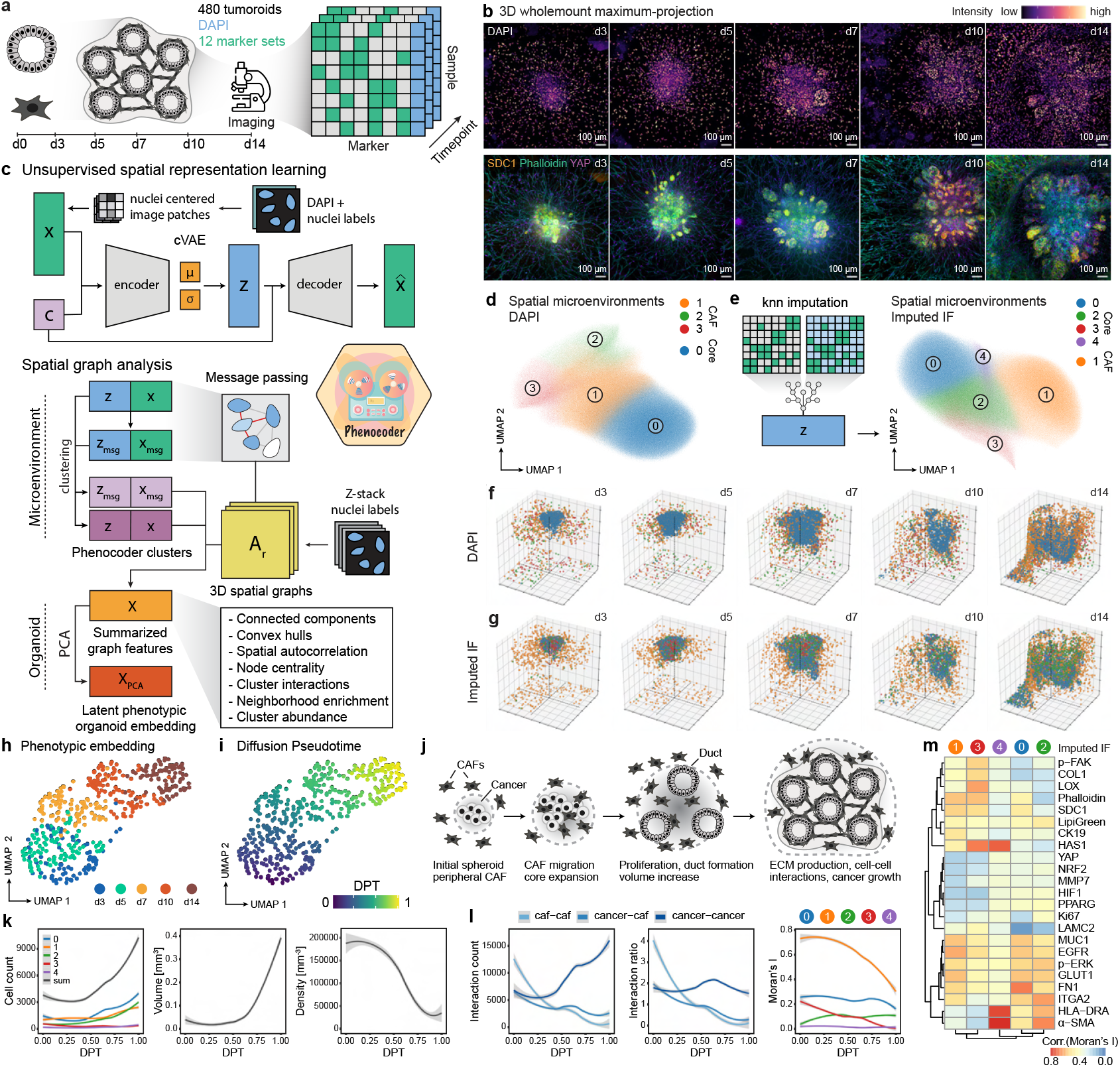
Learning spatiotemporal dynamics in tumoroids with Phenocoder. a) Overview of experimental setup for time-course imaging. Tumoroids (n=480) were cultured for 3, 5, 7,10 and 14 days and subsequently fixed, cleared, stained with 12 different marker sets and imaged. b) Time-course imaging of tumoroid growth using DAPI, SDC1, Phalloidin, and YAP1 staining. c) Network architecture of the conditional variational autoencoder employed with Phenocoder which is designed for learning latent spatial representations of nuclei centered image patches, with plate and z-position of multichannel image patches as conditional inputs. Spatial graph analysis allows for quantifying phenotypic changes through unsupervised clustering, spatial graph feature extraction, and dimensionality reduction to generate latent phenotypic organoid embeddings. d) UMAP of learned latent space of spatial microenvironments based on DAPI stainings. e) Knn-imputation of nuclei marker intensities based on latent spatial representation of DAPI microenvironments, illustrated as UMAP. f,g) 3D spatial visualization of encoded nuclei colored by Phenocoder cluster identity for DAPI based (f) and imputed immunofluorescence (g) shown in cut-open perspectives to reveal internal tumoroid architecture. h,i) UMAP of latent phenotypic organoid embedding, colored by timepoint (h) and diffusion pseudotime (DPT) (i). j) Schematic illustrating spatiotemporal dynamics in tumoroids, showing progression from initial spheroid formation through peripheral CAF positioning, CAF migration toward the core, proliferation and duct formation with volume increase, to final ECM production, established cell-cell interactions, and cancer growth. k,l) Line plots visualizing tumoroid features including cell count, volume and density (k), cell-cell interaction terms and spatial autocorrelation (Moran’s I) values at a radius of 150 pixel (l) along pseudotemporal trajectory. m) Heatmap illustrates correlation of Moran’s I for protein markers and Phenocoder cluster identities.

Following image processing, nuclei segmentation, and feature extraction, we trained a cVAE on nuclei-centered DAPI image patches to learn spatial cellular arrangements within the tumoroid microenvironment while correcting for covariates including batch effects and z-axis photobleaching (Fig. 2c, Fig. 4b,c,g). Unsupervised clustering resolved peripheral CAF networks, ductal structures, and dense fibrotic core regions (Fig. 2d,f, Fig. 4d). We extended the initial latent and immunofluorescence features through spatial message-passing, expanding nuclei-centered environments to broader neighborhood representations (Fig. 2c). Subsequently, we imputed immunofluorescence intensities for both nuclei-based and spatial neighborhood-based features across the complete dataset for all markers by constructing k-nearest neighbor search trees in the cVAE latent space and applied unsupervised clustering to both feature sets (Fig. 2e, Fig. 4e,f,h-j). The integrated multi-marker representation revealed greater tissue heterogeneity within the tumoroid core and at ductal structure interfaces compared to the nuclei-only representation alone (Fig. 2f,g).

To generate organoid embeddings, spatial graph statistics were generated for all cluster sets at different spatial kernel sizes (Fig. 2c). We calculated interaction scores, neighborhood enrichments, cluster abundances, spatial autocorrelation, and supplemented these with descriptive morphological features including duct abundance, duct size, distance of duct to organoid center and organoid volume, to generate a high-dimensional multivariate feature space. Dimensionality reduction produced a phenotypic organoid embedding (Fig. 2h) that enabled reconstruction of spatiotemporal dynamics in tumoroid co-cultures through diffusion analysis and pseudo-temporal trajectory construction (Fig. 2i).

Initially, tumoroids formed dense spheroidal core structures at the well center, surrounded by CAF networks characterized by high cellular density and elevated spatial autocorrelation values for CAF and tumoroid core-associated marker proteins and clusters (Fig. 2j- l, Fig. 4k). Duct formation and cancer cell proliferation led to increased cancer-cancer interactions along the trajectory, while both cancer-CAF and CAF-CAF interactions progressively decreased before stabilizing toward the end of the trajectory (Fig. 2k,l). This stabilized state, characterized by high cancer-cancer interactions and reduced heterotypic cell interactions, creates a microenvironment that promotes sustained cancer growth.

This transition was accompanied by reduced spatial autocorrelation values for CAF and tumoroid core markers (Fig. 2l,m). Together, these dynamic changes reveal self-organization into a steady state of cell-cell and cell-ECM interactions that sustains continuous tumoroid growth, demonstrating that Phenocoder successfully captures the emergence of organized tissue architecture from initially homogeneous co-cultures.

### Learning the tumoroid phenotypic space to assess perturbation effects

We next applied the tumoroid platform and Phenocoder for systematic drug screening. To optimize conditions in a pilot screen, we cultured tumoroids for 14 days and treated them with 14 compounds (Table S3) from days 4, 7, and 11 at 1, 5, and 10 µM (Fig. 3a). The compound panel included targeted inhibitors (receptor tyrosine kinases, intracellular signaling, cell adhesion, metabolism, and ECM modification), standard-of-care chemotherapeutics (e.g. Gemcitabine, Paclitaxel), positive controls (SN38, Bortezomib), and DMSO controls. We acquired imaging data at 20x resolution for 20 µm z-stacks for two imaging cycles (cycle 1: DAPI, SDC1, ITGA2, and LAMC2; cycle 2: DAPI, COL1, KI67, and CK19), including an elution cycle to validate multiplexing consistency (Extended Data Fig. 5a). After quality control, we extracted intensity and morphological feature sets for ∼ 4.2 million segmented nuclei from 741 imaged tumoroids and applied Phenocoder to encode spatial cellular arrangements based on multichannel nuclei-centered image patches for each imaging cycle (Fig. 3b). Subsequently, a latent phenotypic tumoroid embedding was constructed to quantify drug perturbation effects.

**Figure 3.**
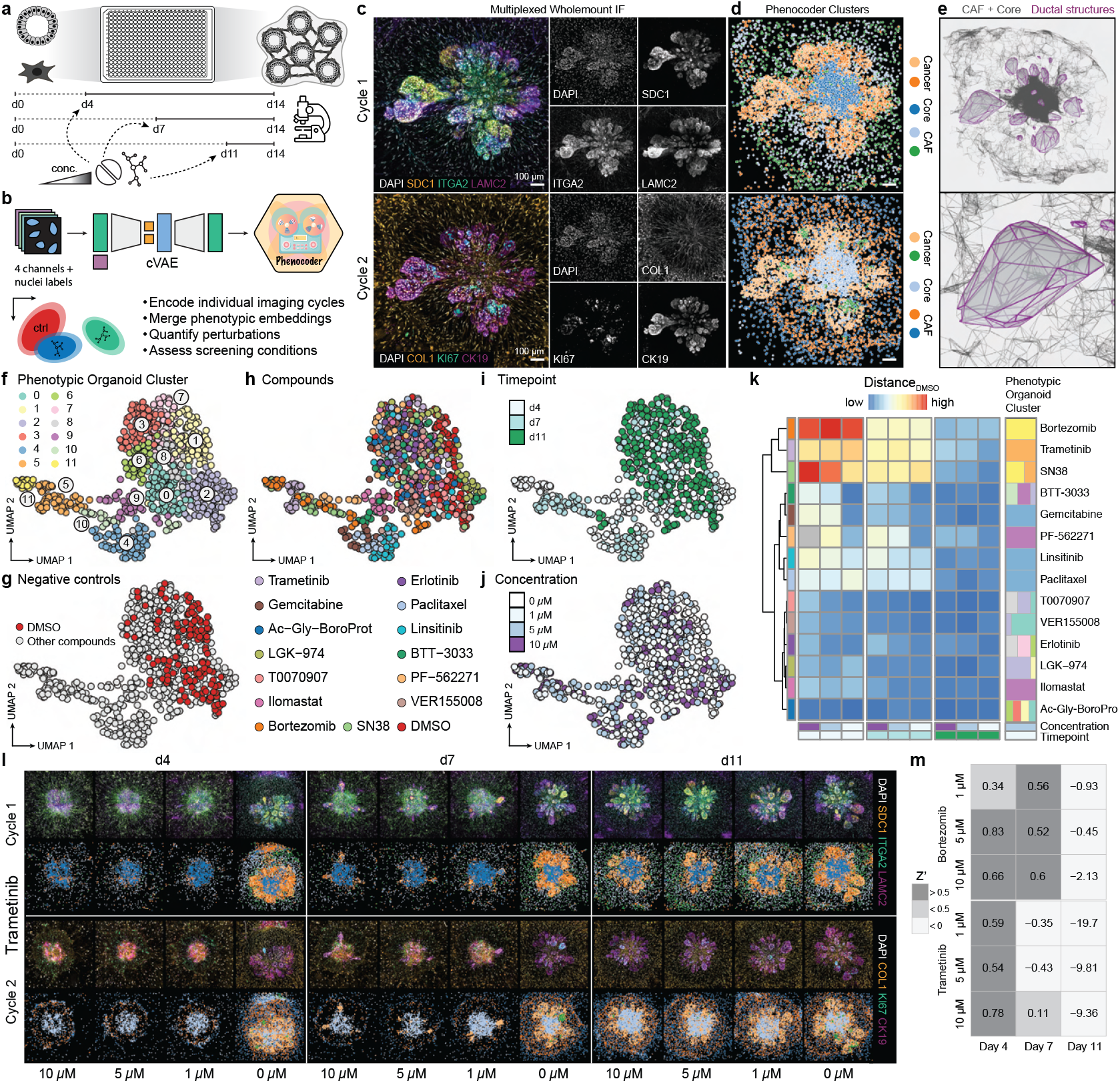
Learning perturbation effects in the tumoroid phenotypic space. a) Experimental design overview showing tumoroid culture timeline (day 0–14) with drug perturbation design for high content screening: 14 compounds are tested at three concentrations (1, 5, and 10 µM), treated from three distinct timepoints (d4, d7, or d11) until day 14, including DMSO controls, and SN38 and Bortezomib as positive controls. b) Phenocoder analysis workflow process for pilot screen data. c) Representative multiplexed whole-mount immunofluorescence images for two imaging cycles. DAPI (white), cycle 1: SDC1 (orange), ITGA2 (green), LAMC2 (magenta), and cycle 2: COL1 (orange), Ki67 (green), CK19 (magenta). Right subpanels: grayscale images of individual channels. d) Spatial visualization of Phenocoder-derived clusters for both imaging cycles, revealing distinct microenvironmental organization of cancer and fibroblast populations, segmenting ductal regions, dense fibrotic core, and fibroblast network. e) Rendering of fibroblast network and tumoroid core as spatial graph (gray) with segmented connected components of ductal cancer structures (magenta). f-j) UMAP of latent phenotypic organoid embedding, colored by organoid cluster (f), DMSO controls (g), compounds (h), treatment timepoint (i) and compound concentration (j). k) Heatmap quantifying treatment effects via grouped Mahalanobis distances to DMSO control across compounds, timepoints, and concentrations. Side bar shows the phenotypic organoid cluster (f) composition for each compound at treatment condition day 4 and concentration 5 µM. l) Overview montage of Trametinib treatment effects showing original immunofluorescence (top rows) and corresponding Phenocoder clusters (bottom rows) across all timepoints and concentrations. m) Statistical validation of assay quality using Z’ factor values for Bortezomib (positive control) and Trametinib across treatment conditions.

**Figure 4.**
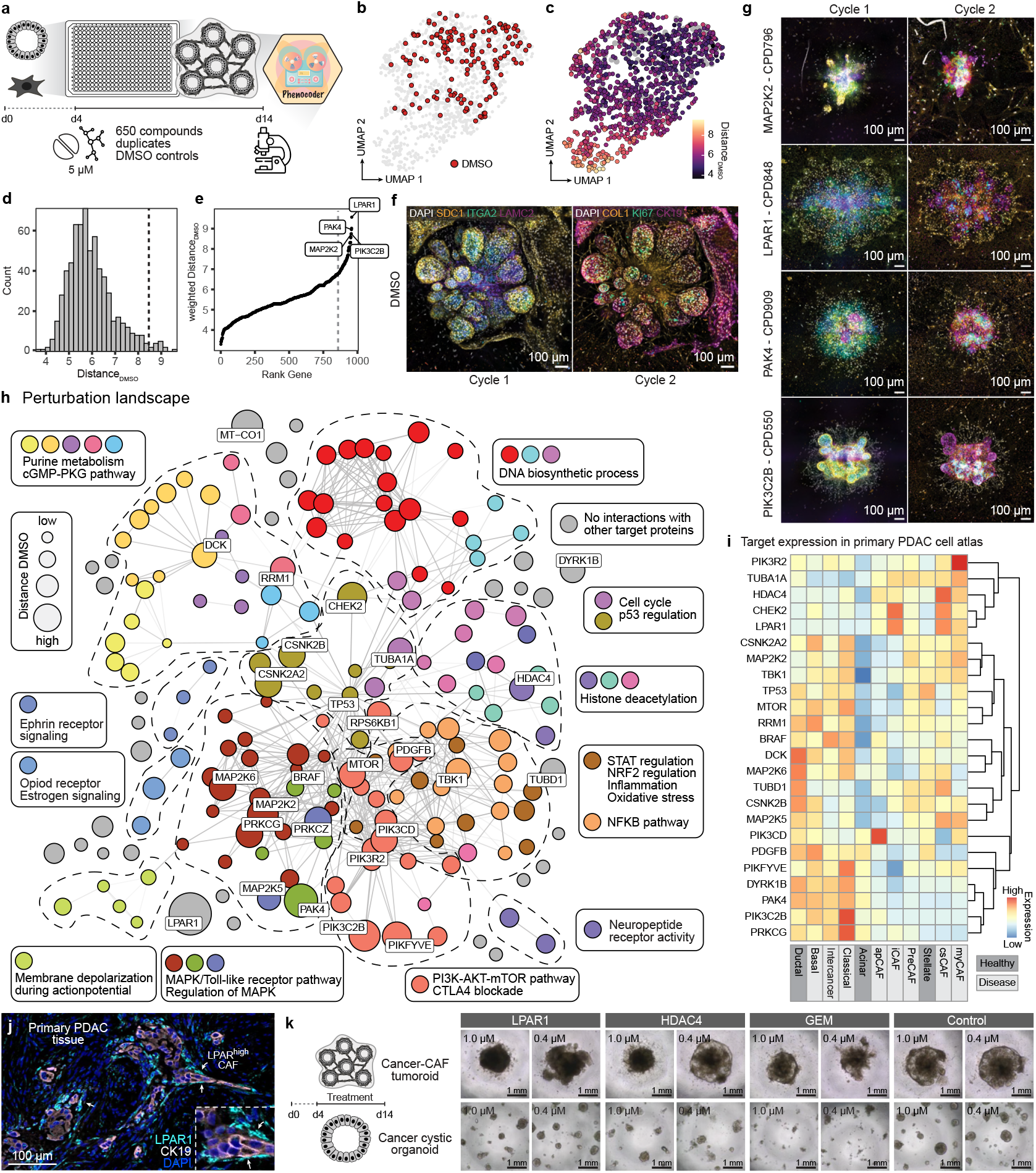
Systematic perturbation screening of cancer-CAF interactions. a) Experimental design of high-content screening (HCS) for 650 compounds in tumoroid cultures for 14 days, with treatment applied from day 4 onwards at 5 µM followed by multiplexed whole-mount imaging and drug effect assessment through Phenocoder. b-c) UMAP embeddings of organoid phenotypic features showing the distribution of DMSO controls (b, red) and the phenotypic effects of compounds colored by distance from DMSO controls (c). d) Histogram shows distribution of compound effect sizes (DistanceDMSO). e) Weighted distance ranking of targeted genes with top four hits (LPAR1, MAP2K2, PIK3C2B and PAK4) highlighted. f) Representative immunofluorescence images of PDAC tumoroid cultures under DMSO condition across two imaging cycles. DAPI (white), cycle 1: SDC1 (orange), ITGA2 (green), LAMC2 (magenta), and cycle 2: COL1 (orange), Ki67 (green), CK19 (magenta). g) Representative images showing phenotypic effects of compounds targeting MAP2K2, LPAR1, PAK4, and PIK3C2B across two imaging cycles with same multiplexed marker staining as in (f). h) Perturbation landscape of tumoroid co-cultures represented as a STRING interaction network of the top 100 target proteins ranked by weighted DMSO distance, with labeled targets of the 16 most effective compounds. Nodes are color-coded by biological functions, with node size proportional to phenotypic effect. i) Heatmap showing the expression of top compound targets from the screen in the integrated PDAC scRNA-seq atlas (Xu et al., a) across cancer and fibroblast subpopulations. j) Immunohistochemistry of primary PDAC tissue highlighting LPAR1 expression (cyan) on CAFs in proximity to cancer cells (CK19, white). k) Brightfield images showing follow-up validation of selected compounds and DMSO control in tumoroids co-cultures (top) or cancer cystic organoid cultures (bottom). LPAR1 and HDAC4 inhibitors show strong effects only on the tumoroid and have limited effect on cystic cancer organoids.

The autoencoder demonstrated robust training and high reconstruction fidelity (Extended Data Fig. 5b,e,f), preserving enhanced spatial information compared to conventional single-cell immunofluorescence and morphology-based embeddings (Fig. 5c,d). Multiplexed imaging data for DMSO controls revealed robust and distinctive marker patterns (Fig. 3c, Extended Data Fig. 7a). Unsupervised clustering in the latent space identified 3D cellular arrangements in tumoroids across imaging cycles (Fig. 3d, Extended Data Fig. 6a-d,Extended Data Fig. 7b). The first imaging cycle segmented ductal cancer structures (clusters 2, 3) with distinctive LAMC2^+^ zones (cluster 3), dense fibrotic core (cluster 0), and a peripheral CAF network (clusters 1, 4) (Fig. 6a,b,e). The second imaging cycle similarly detected ductal cancer structures (clusters 3, 4) while revealing high proliferative zones (cluster 4, KI67^+^), a dense fibrotic core (cluster 1), and a peripheral CAF network (clusters 0, 2) (Fig. 6c-e). Clustering remained consistent across imaged plates and drug perturbation conditions (Extended Data Fig. 7b).

**Figure 5.**
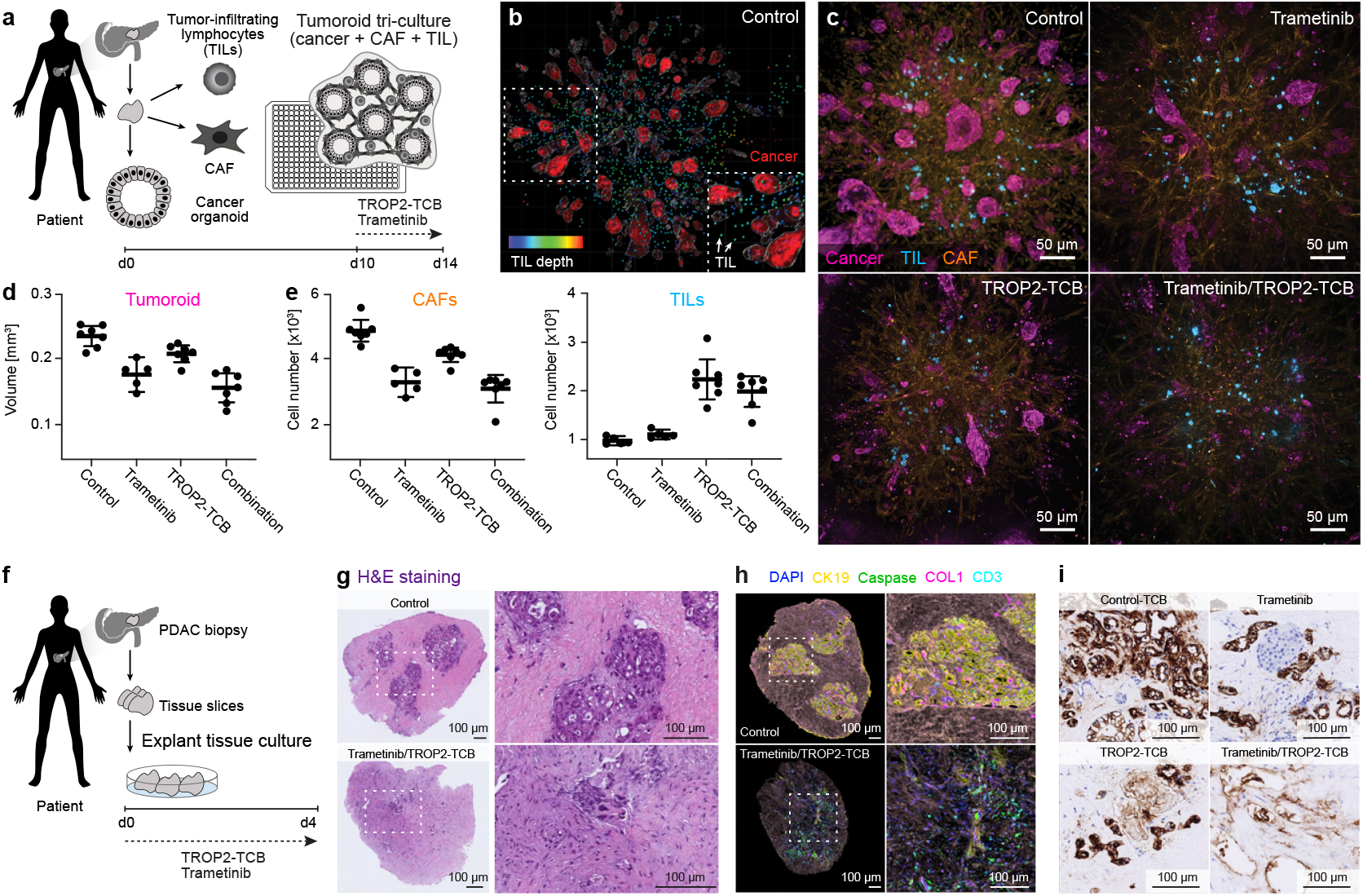
Combinatorial MEK inhibition and T cell bispecific antibody treatment induces cancer cell death in multilineage tumoroids and PDAC explants. a) Overview of the high-content imaging-based phenotypic screening platform for patient-derived cancer-CAF tri-culture PDAC tumoroids including tumor infiltrating lymphocytes (TILs) or activated T cells from peripheral blood mononuclear cells (PBMCs). b) 3D rendering of whole-mount imaging of cleared tri-culture tumoroids stained for cancer cells (CK19) and TILs (CellTrace™ Far Red). c) Maximum projection of 3D whole-mount imaging tri-culture tumoroids for control condition, treated with Trametinib, TROP2 T cell bi-specific antibody, or combination of Trametinib and TROP2-TCB. Tumoroids stained for cancer cells (CK19), CAFs (Collagen I), and TILs (CellTrace™ Far Red). d,e) Quantification of tri-culture tumoroid volume and CAF/TIL cell counts across treatment groups. f) Overview of explant tissue culture from patient-derived PDAC biopsies. g-i) Imaging of explants with H&E staining (g), immunofluorescence staining for CK19, cleaved Caspase-3, Ki67, CD3, and Collagen I with DAPI as a nuclear counterstain (h), and immunohistochemistry for pERK (i) in PDAC explant cultures under each treatment condition.

In addition, this clustering approach allowed segmentation of ductal structures as connected components on spatial graph representations at the individual tumoroid level, visualized as 3D renderings of CAF networks and segmented ductal structures (Fig. 3e).

The latent phenotypic organoid embedding facilitated identification of phenotypic organoid clusters (Fig. 3f, Fig. 6f) and enabled detection of treatment duration and concentration-dependent effects (Fig. 3g-j). For each treatment, we estimated effect size by calculating grouped Mahalanobis distance, revealing that treatment timepoint exerted the strongest impact on perturbation effects, followed by compound concentration (Fig. 3k). Phenotypic cluster proportions shifted with treatment conditions and displayed compound specificity (Fig. 3k, Extended Data Fig. 6f). These phenotypic clusters captured the heterogeneity across multivariate spatial graph-based features including ductal architecture metrics, cell-cell interaction patterns, and marker expression ratios, demonstrating that the observed organoid phenotypes reflect coordinated changes in tissue organization and cellular composition (Extended Data Fig. 6g).

Trametinib, an inhibitor of the mitogen-activated protein kinase enzyme MEK2 (MAP2K2), demonstrated comparable effect sizes to positive controls Bortezomib and SN38 (Fig. 3k,l). Linear discriminant analysis in the latent phenotypic organoid space yielded Z’ prime values exceeding 0.5 (Fig. 3m) at treatment timepoints day 4 and 7, validating high assay quality (Bar and Zweifach 2020). Treatment starting at day 4 at 5 µM was selected for further screening, as 10 µM samples showed susceptibility to detachment after the elution cycle (Extended Data Fig. 7a). This pilot screen demonstrated that Phenocoder enables systematic evaluation of screening conditions and provides a robust computational framework for quantifying drug-induced phenotypic changes in heterogeneous 3D multicellular systems.

### High-throughput perturbations of cancer-CAF interactions

To systematically characterize the perturbation landscape of cancer-CAF interactions in PDAC tumoroids, we conducted a large-scale phenotypic screen with 650 compounds (Table S4). Compound selection was guided by differential gene expression and pathway enrichment analysis of tumoroids profiled over time, benchmarked against scRNA-seq data from primary PDAC biopsies and liver metastases to align with in vivo transcriptional states (Xu et al., a). We applied the optimized screening conditions (day 4, 5 µM) determined in the pilot screen and analyzed the acquired imaging data (∼ 12TB) with Phenocoder (Fig. 4a). We observed consistent multiplexing across imaging cycles (Extended Data Fig. 8a) and the autoencoder demonstrated stable training convergence with high fidelity image reconstruction (Extended Data Fig. 8b,c). After quality control, we encoded ∼ 12 million spatial microenvironments across 1391 tumoroids and unsupervised clustering revealed distinct microenvironmental domains that consistently segmented ductal structures, peripheral spreading regions, dense fibrotic cores, and CAF networks in both imaging cycles (Extended Data Fig. 9a-g).

In the phenotypic organoid embedding space (Fig. 4b), we quantified DMSO control variability and calculated effect sizes for each screened compound (Fig. 4c,d). We aggregated perturbation effects at gene-target level (Fig. 4e), and mapped them to a STRING-based protein–protein interaction network (Szklarczyk et al.)(Fig. 4h), enabling enrichment analysis of biological pathways and construction of a perturbation landscape that highlights the druggability of cancer-CAF interactions in tumoroids. Our analysis identified several highly druggable pathways including MAPK, mTOR, and NF-kappa signaling networks. Top hits included LPAR1, PAK4, PIK3C2B, PIKfyve and MAP2K2, which exhibited significant phenotypic changes compared to DMSO controls (Fig. 4f,g). LPAR1 (Lysophosphatidic acid receptor 1) mediates LPA signaling, promoting CAF activation and extracellular matrix remodeling, processes that contribute to fibrosis-associated tumor progression in PDAC (Volat et al.). Additionally, PAK4, a regulator of cytoskeletal dynamics, has been implicated in enhancing fibroblast migration and proliferation in response to tumor cell signaling, potentially promoting cancer progression (Li et al., a; Xu et al., b). PIKfyve was identified as a metabolic vulnerability in PDAC, where its inhibition disrupts lysosomal lipid homeostasis and synergizes with KRAS-MAPK blockade (Cheng et al., a). PIKfyve inhibition has also been linked to fibrosis regulation in cardiac and pulmonary models (Barkovskaya et al.; Cinato et al.).

Furthermore, MAP2K2, a key component of the MAPK pathway, is involved in the fibroblast-mediated response to stress and growth signals within the tumor microenvironment, and its inhibition has shown promise in disrupting both fibrosis and cancer-associated fibroblast (CAF) signaling (Nguyen et al.; Caunt et al.). Notably, the identification of MAP2K2 was consistent with the strong Trametinib effect from the pilot screen. The top compounds induced distinct organoid phenotypes (Extended Data Fig. 9g), which displayed considerable phenotypic heterogeneity that is linked to target gene expression in primary PDAC tissue (Fig. 4i, Extended Data Fig. 9h,i). These pathway level effects reflect hallmarks of PDAC biology, including desmoplasia, tumor progression, and resistance to standard therapy, highlighting that synergy between cancer cells and CAFs is a key driver of the observed phenotypes.

We focused on the target LPAR1 for follow-up evaluation. LPAR1 is expressed in multiple CAF states, with the highest transcript detection in myCAF, iCAF and csCAF (Fig. 4i). IHC revealed that this LPAR-high CAF population is located directly adjacent to CK19^+^ cancer cells within primary PDAC tissue (Fig. 4j). We treated cystic cancer organoids and cancer-CAF tumoroids with the LPAR1 inhibitor, Gemcitabine and an HDAC4 inhibitor, and found that LPAR1 had strong effect on the tumoroid but limited effect on the cancer organoids in the absence of CAF. Interestingly, HDAC4 inhibition showed a similar CAF-dependent phenotype, whereas Gemcitabine induced cancer cell death in both contexts. LPAR1 is a target that has been used in the treatment of pulmonary fibrosis (Volkmann et al.), and our data suggests that it could be a novel selective target of the PDAC stromal microenvironment. Notably, LPAR1 can signal through the MEK/ERK pathway, and several other compounds targeting this pathway resulted in disrupted tumoroid growth. Altogether, these data reveal that unsupervised machine learning based on high-throughput, high information content phenotyping data of complex multicellular cancer microenvironments in vitro can identify pathways and targets that would not be discovered in standard cancer cystic mono-cultures relying on singular readouts such as cell death or organoid volume.

### CAF disruption enables immune targeting of cancer cells

T cell-bispecific (TCB) antibodies are an emerging class of immunotherapies that recruit polyclonal T cells toward tumor-associated antigens, representing a key advance in precision oncology (Klein et al.; Herrera et al.). We focused on TROP2 as a target given its broad expression across epithelial cancers and clinical validation. Notably, antibodies-drug conjugates (ADCs) directed against TROP2 have demonstrated significant benefit in patients with advanced solid tumors, with progression-free and overall survival significantly improved in clinical trials (Bardia et al.). On this basis, we evaluated a TROP2-directed TCB in our tri-culture tumoroid platform, consisting of patient-derived cancer cells, matched CAFs, and T cells, to assess combinatorial strategies that integrate pathway inhibition with immune cell engagement (Fig. 5a, Extended Data Fig. 10a). In PBMC tri-cultures, treatment with Osimertinib (EGFR inhibitor) modestly reduced cancer cells outgrowth, but this effect was limited and not sufficient to induce strong immune activity. Addition of a control TCB antibody produced no benefit, whereas combining Osimertinib with a TROP2-TCB enhanced T cells infiltration and triggered detectable cancer cell death (Extended Data Fig. 10b-d). Perturbations at multiple levels of the MAPK pathway disrupted tumoroid growth in our screen, and CAF states linked to PDAC severity have enhanced MEK/ERK activity. We treated tri-culture tumor-infiltrating lymphocyte (TIL) tumoroids with Trametinib (MEK inhibitor), and a T cell-bispecific antibody targeting TROP2, either alone or in combination, from day 10 to 14 of culture. We then compared them with control tumoroids (Fig. 5a-c, Extended Data Fig. 10e-j). As previously observed in our experiments, Trametinib resulted in disruption of the fibroblast network, overall reducing tumoroid growth and proliferation (Fig. 5c,d). TROP2-TCB treatment induced T cell proliferation and cancer targeting, and combined with Trametinib led to enhanced cancer cell death (Fig. 5d,e).

To validate these results, we established patient-derived PDAC explant cultures that preserved tissue integrity, including ductal structures, stromal fibroblasts, and infiltrating immune compartments, for up to 4 days in culture (Fig. 5f). This model enabled assessment of therapeutic responses in a setting that maintains the native tumor architecture, spatial organization, and direct cell–cell interactions between cancer, stroma, and immune compartments. Combination treatment with Trametinib and TROP2-TCB induced marked remodeling of the explants, characterized by loss of organized fibroblastic domains, expansion of TILs within stromal and epithelial regions, and widespread tumor cell clearance (Fig. 5g-i). Histological and immunofluorescence analyzes corroborated these effects, showing reduced Ki67^+^ proliferating cells, increased caspase-mediated apoptosis, and progressive collapse of ductal epithelial structures. In addition, regions of dense fibrotic stroma were replaced by infiltrating immune cells, indicating that the combination regimen not only disrupted tumor growth but also reshaped the microenvironment to favor immune-mediated elimination.

## Discussion

Quantitative spatial phenotyping of multilineage organoid systems addresses a fundamental bottleneck in modeling disease microenvironments. We established an integrated experimental-computational platform combining modular tumoroid culture with machine learning-based spatial analysis, demonstrating its capabilities in PDAC tumoroids incorporating stromal and immune components. We applied the framework to study spatiotemporal phenotypic dynamics, systematic drug perturbation response, and immunotherapy targeting in tumoroid co-cultures.

We developed Phenocoder, a machine learning framework that combines conditional variational autoencoders with spatial graph analysis to generate latent phenotypic embeddings of complex tissue architectures. Phenocoder extends existing computational approaches by integrating both cell-intrinsic and microenvironmental features across multiple spatial scales, enabling robust and quantitative assessment of cellular organization and cancer-CAF interactions in heterogeneous 3D tumoroid structures. This framework addresses technical challenges specific to 3D co-culture systems, including intrinsic morphological variability between tumoroids, z-axis photobleaching in thick specimens, and batch effects between experimental plates, enabling reliable and reproducible analysis of our high-throughput cancer-CAF interaction screen. The generalizable analytical framework established here for learning multivariate feature tumoroid profiles can be extended to other cancer types with prominent stromal components and is broadly applicable to any organoid system, offering a scalable platform for development of microenvironment-directed therapies. The platform also provides a versatile method for testing immunotherapy combinations with stroma- or tumor-targeting therapies, by integrating the appropriate immune cells.

We identified established and previously underappreciated druggable targets in the PDAC microenvironment. MAP2K2 inhibition further confirmed the relevance of the well studied MAPK pathway in PDAC progression. LPAR1 is notable given its established role in fibrosis with respective inhibitors in clinical trials for pulmonary disease (Corte et al.) marking potential for drug repurposing to selectively target stromal remodeling in PDAC. Lysophosphatidic acid is a mitogen and has extensive roles in cancer biology, and LPAR1 has been linked to PDAC metastasis (Juin et al.). PIKfyve, recently identified as a metabolic vulnerability in PDAC where its inhibition disrupts lysosomal lipid homeostasis and synergizes with KRAS-MAPK blockade (Cheng et al., a), also emerged from our analysis. PIKfyve inhibitor (ESK981) is currently in clinical evaluation in solid tumors, including PDAC, indicating its relevance for clinical translation.

Integration of phenotypic screening data with an integrated PDAC single-cell transcriptome atlas can enable rational selection of therapeutic candidates based on expression patterns and patient prevalence in specific cellular compartments of primary tumors. This strategy provides a valuable framework for validating target expression across diverse cell subpopulations, offering insights into which cellular compartments might be affected by specific compounds. Such cell-type specific targeting potential offers a pathway toward precision medicine approaches in PDAC, where treatment strategies could be tailored to universal fibroblast and cancer subtypes, as well as states that stratify individual patients.

Despite these advances, several limitations remain. The tumoroid model lacks key components of the native PDAC microenvironment including vasculature, neurons, and immune populations, particularly tumor-associated macrophages and T cells that engage in crosstalk with CAFs to establish immunosuppressive niches (Halbrook et al.; Raffo-Romero et al.; Palikuqi et al.). The absence of these components limits the ability to assess how stromal-targeted therapies might influence anti-tumor immunity, an important consideration as immunotherapy combinations are explored in PDAC. Furthermore, while the phenotypic organoid embeddings effectively capture treatment-induced architectural changes, deeper analysis connecting specific features to drug mechanisms of action and underlying protein interaction networks would further enhance interpretability (Ramos Zapatero et al.). Technical challenges include the inability of the system to fully recapitulate the dense stromal barriers that impede drug delivery in vivo. Additionally, compound screening performed on a single patient-derived tumoroid line requires validation across multiple patient backgrounds to address inter-tumor heterogeneity in phenotypes and drug sensitivities.

This integrated experimental-computational framework establishes a generalizable and scalable approach for spatial phenotyping of multilineage organoid systems where cell-cell interactions drive disease biology, providing a powerful platform for identifying therapeutic vulnerabilities such as those at the cancer-stroma interface in PDAC. By systematically mapping the phenotypic consequences of targeted perturbations in a biomimetic tumoroid system, we reveal promising opportunities for disrupting the pathological relationships between cancer cells and fibroblasts that drive disease progression. As similar stromal barriers limit therapeutic efficacy across multiple solid tumors, this approach offers a blueprint for developing microenvironment-directed treatment strategies that could improve outcomes not only in pancreatic cancer but potentially in other fibrosis-associated malignancies.

## Data availability

Sequencing data will be made available upon publication. Imaging data is available upon request to the corresponding authors.

## Code availability and analytic reproducibility

Phenocoder and other codes used in the analyses are available upon publication on GitHub under: https://github.com/devsystemslab/pdac_tumoroid

## Author contributions

RO performed IHC on primary PDAC tissue. MS and IC performed IHC on tumoroid samples. RO established and banked the organoids and fibroblasts lines from biopsy tissues. RO and HM generated and expanded organoids and fibroblasts used in this study. RO established, designed and performed the automated multiplexed whole-mount HCS experiments with support from IL and HM and advice from MB. CH developed the computational pipeline for whole-mount HCS image data processing and implemented the Phenocoder algorithms. CH processed and analyzed wholemount imaging data with support from PS. QX performed computational analyses to compare screening data with the integrated single cell PDAC atlas. BG, QX and IL informed and designed the compound library. RO and LS established immune cell co-culture systems with support from LC. RO and EA designed and performed explant culture systems with support from MPL. CH, RO and JGC designed the study, interpreted results and wrote the manuscript.

## Acknowledgements

We acknowledge the support of the non-profit foundation HTCR, which holds human tissue on trust, making it broadly available for research on an ethical and legal basis. We thank Elizabeth Pando Rau from the Hepatobiliopancreatic Surgery and Adult Liver Transplant Service at Vall d’Hebron University Hospital (Spain), and BeCytes Biotechnologies (Spain) for facilitating access to biospecimens and ensuring their availability for research in compliance with ethical and legal standards, following the provision of informed consent by the donor. We are grateful to Isabell Rall and Laura Gaspa Toneu for laboratory management support, and the IHB Biobank team for sample processing assistance.

## Methods

### Human samples

Human PDAC tissue samples and associated data were obtained (Table S1), and all experimental procedures were performed within the framework of the non-profit foundations HTCR (Munich, Germany) and Cyte (Barcelona, Spain), as well as the University of Basel (Basel, Switzerland). Informed consent was obtained from all patients prior to sample collection.

The HTCR and Cyte Foundations, as well as the University of Basel, operate under ethical approvals from the Ethics Committee of the Faculty of Medicine at LMU (approval number 025-12) and the Bavarian State Medical Association (approval number 11142). Additional approvals were granted by the Hospital Universitari Vall d’Hebron (approval number PR(AG)137/2023, CODE: D0887/22.2023.V1) and the University Hospital Basel (EKNZ approval number 2019-02118).

### PDAC tissue preparation and isolation

The surgical tissue is washed several times with Dulbecco’s phosphate buffered saline (DPBS). The tissue was finely chopped using surgical scissors and a scalpel. The tissue was transferred to a gentleMACS C tube (miltenyo Biotec, 130-093-237) and washed again with DPBS. The washed tissue was digested with the dissociation mix at 37°C for 60 minutes. The dissociation mix was prepared with collagenase type2 (Thermo Fisher Scientific, 10738473, 5 mg/ml), DNAse (Sigma-Aldrich, D5025-150KU, 7µg/ml), Hyaluronidase (Sigma-Aldrich, H6254-500MG, 270 µg/ml), Trypsin inhibitor (Sigma-Aldrich, T6522-100MG, 1 mg/ml), Y-27632 (STEMCELL Technologies, 72307, 10 µM). The tissue pieces were digested with gentleMACS Octo Dissociator program “37c_h_TDK_1” (Miltenyi Biotec, 130-134-029), as per the manufacturer’s instructions. Tissues were enzymatically treated and then washed with DMEM containing 10% fetal bovine serum (Thermo Fisher Scientific, A3160502) to stop the enzymatic reaction. The Cell suspension was then washed with MACS buffer and autoMACS rinsing solution before proceeding with magnetic activated cell sorting (MACS). For the cell isolation, the suspension was incubated with CD326 (EpCAM) MicroBeads (Miltenyi Biotec, 130-061-101, 10 µL beads in 100 µL MACS buffer) for 20 minutes on ice. EpCAM-positive and EpCAM negative cells were further processed for immune and fibroblast cell isolation by incubation with CD45 MicroBeads microbeads (Miltenyi Biotec, 130-045-801, 10 µL beads in 100 µL MACS buffer) for 20 minutes on ice. CD45 positive cells were cultured as tumor infiltrating lymphocytes (TILs), while CD45 negative cells were cultured as cancer associated fibroblasts.

### Establishment of pancreatic cancer organoids

The obtained pancreatic cancer cells were embedded in growth factor reduced (GFR) Matrigel (Corning, 356231) and cultured in the following complete medium. Advanced DMEM/F12 (Thermo Fischer Scientific, 12634028), Primocin® (Invivo-Gen, ant-pm-1, 1 mg/ml), GlutaMAX (Invitrogen, 35050061, 1x), 1x B27( Invitrogen, 17504044, 1x), Gastrin (Millipore, G9145-.1MG, 1 nM), N-acetyl-L-cysteine (Sigma-Aldrich, A9165, 1mM), Nicotinamide (Sigma-Aldrich, 72340-100G, 10mM), A83-01(Tocris, 2939, 0.5 µM), Noggin (Peprotech,120-10C, 0.1 µg/ml), R-Spondin1 (Peprotech, 120-38, 100 ng/ml), Wnt3A (RD, 5036-WN-500/CF, 50 ng/ml), EGF (Peprotech, AF-100-15, 50 ng/ml), FGF10 (Peprotech, 100-26, 100 ng/ml). Y-27632 (STEMCELL Technologies, 72307, 10 µM) was added for only one day after starting the organoid culture, and on the following day, the cells were cultured in a complete medium without Y-27632. The medium was changed 2 to 3 times a week. All lines used in the studies were verified as negative for mycoplasma before further experimentation.

### Establishment of cancer associated fibroblasts

The obtained fibroblast cells were resuspended in mesenchymal stem cell growth medium (MSCGM2, promocell, C-28009) and seeded onto a cell culture plate. The medium was replaced 2–3 times per week. Fibroblasts were cultured under standard conditions at 37°C in a 5% CO_2_ and 20% O_2_. All lines used in the studies were verified as negative for mycoplasma before further experimentation.

### Establishment of tumor infiltrating lymphocytes

Tumor infiltrating lymphocytes (TILs) were seeded in ultra-low attachment (ULA) round-bottom plates (Corning) using ImmunoCult™-XF T Cell Expansion Medium (Stemcell Technologies, 10981) as the base medium. T cells were activated with 25 µL/ml of ImmunoCult™ Human CD3/CD28 T Cell Activator (Stemcell Technologies, 10971) and incubated at 37°C in a humidified incubator with 5% CO_2_ for 3 days. Following activation, cells were harvested, centrifuged at 320g for 5 min at 4°C, and resuspended in ImmunoCult™-XF T Cell Expansion Medium supplemented with 100 IU/ml recombinant human IL 2 (PeproTech). CD3/CD28 stimulation was omitted at this stage. Cells were counted using a hemocytometer or automated cell counter (Thermo Fisher Countess™II) and reseeded in cytokine-containing medium for an additional 4 day resting period. Purity of the enriched cells was validated by flow cytometry. All lines used in the studies were verified as negative for mycoplasma before further experimentation.

### Pancreatic cancer organoid expansion culture

PDAC organoids are initially expanded in 6-well plate dome cultures before being transferred to the CERO bioreactor for scale-up to facilitate high-throughput screening. Each well is washed with DPBS, and domes are scraped and resuspended in TryPLE Express (Thermo Fisher Scientific, 10718463) containing 10 µM Rock inhibitor. Organoids are dissociated by pipetting, transferred to a 50 ml Falcon tube, and incubated on a rocker at 37°C for 8–10 minutes. The reaction is stopped with warm DMEM++, followed by centrifugation at 300g for 5 minutes at 4°C. The supernatant is aspirated, and the pellet is resuspended in warm DMEM++, washed once more, and mixed with Cultrex Type 2 (R&D Systems, 3533-005-02) while kept on ice. Cells are plated as 35 µl domes in a 6-well plate, incubated at 37°C for 20–60 minutes, and supplemented with 2 ml of complete PDAC medium containing 10 µM Rock inhibitor. The medium is changed every 3 days to support organoid growth. Following expansion, organoids are transferred to the CERO bioreactor for large-scale culture and phenotypic screening. Organoids are cultured in the bioreactor in complete medium supplemented with 5% Cultrex, and the medium is changed once a week. If the medium color began turning yellow before the scheduled change, fresh medium was added to maintain optimal growth conditions.

### Fibroblast expansion culture

Fibroblasts are maintained and passed upon reaching 70% confluency, the medium is aspirated and cells are washed with DPBS, TryPLE Express is added, and cells are incubated at 37°C for 3 minutes. Once detached, the reaction is stopped with cultured medium. The suspension is centrifuged at 300g for 3 minutes, washed, and counted. 2.5×105 cells are resuspended in culture medium and plated in T175 flasks, reaching confluency within 6 days.

### Tumor infiltrating lymphocyte culture

Established tumor infiltrating lymphocytes were collected exclusively from the bottom of each U bottom well, washed once with base medium, and centrifuged at 320g for 5 min at 4°C. Cells were counted, and 0.1–1×10^6^ cells were seeded per well in 96-well U-bottom plates (Corning) containing ImmunoCult™-XF T Cell Expansion Medium supplemented with 100 IU/ml recombinant human IL-2. This process of harvesting, washing, centrifugation, counting, and reseeding was repeated as necessary until the desired cell numbers were achieved.

### Co-culture of tumoroid with cancer cells and CAFs

Tumoroids were co-cultured with pancreatic cancer cells and cancer-associated fibroblasts (CAFs) to establish a physiologically relevant in vitro model. Cancer cells and CAFs were dissociated using TrypLE Express (TrypLE EX), as described in pancreatic cancer organoid and fibroblast culture protocols. A 384-well plate was coated with a 60% Cultrex Type 2(R&D Systems) diluted in culture medium (DMEM/F-12+++ and EGM-2). The coating process was performed using an automated liquid-handling system (Integra AssistPlus) to ensure consistency and reproducibility. Following gel coating, dissociated cancer cells and CAFs were mixed with culture medium supplemented with Y-27632 (10 µM) to enhance cell survival. The cell suspension was then seeded onto the coated gel layer using the automated system (Intera ViaFlo). After seeding, the co-cultures were maintained under standard conditions at 37°C in a humidified incubator with 5% CO_2_. The medium was replaced daily using an automated liquid handler (Biotek EL406) to ensure consistent nutrient supply and dead cell removal. The co-culture of cancer cells and CAFs was maintained for 14 days.

### Tri-culture of tumoroid with cancer cells, CAFs and T cells (TILs or PBMC-derived activated T cells)

Expanded TILs or PBMC-derived activated T cells were tri-cultured with cancer cells and CAFs using the same automated culture program. T cells were stained with the CellTrace™ Far Red (Thermo Fisher, C34564) following the manufacturer’s instructions, using a final working concentration of 1 µM. The tri-cultured tumoroid system utilized a combination of RPMI 1640 (Thermo Fisher Scientific, 61870127), DMEM/F-12+++, and EGM-2 (Lonza, CC-3162) supplemented with 100 IU/ml recombinant human IL-2 as the culture medium to support optimal growth and interaction of the different cell types. The coating process for the tri-culture was also performed using the automated liquid-handling system (Integra AssistPlus, ViaFlo, and Microplate Washer Dispenser, BioTek EL406) to ensure consistency and reproducibility. The tri-culture was maintained for 11 days under the same culture conditions, with daily medium replacement to sustain cell viability and functionality.

### Tumoroid 3D clearing wholemount staining

Samples from the co-cultured and tri-cultured tumoroid model were collected at multiple time points and fixed with 4% paraformaldehyde (PFA) at room temperature for 45 minutes. Following fixation, samples were washed thoroughly with phosphate buffered saline (PBS) to remove residual PFA. For permeabilization, samples were incubated overnight at 4°C in a permeabilization buffer consisting of 4% Tween-10 and 1% triton X-100 in PBS. This was followed by an overnight incubation at 4°C in a blocking buffer containing 6% donkey serum, 1% bovine serum albumin (BSA), 4% Tween-20, and 1% Triton X-100 to minimize nonspecific binding. Primary and secondary antibodies were prepared in the blocking buffer and incubated with the samples for 2–3 days at 4°C to ensure thorough penetration of the 3D structures. After antibody incubation, samples were washed extensively with PBS to remove free antibodies. The samples were then transferred into a fructose-base optical clearing buffer to enhance image clarity. Once the clearing process was complete, image acquisition was performed. All steps, including fixation, permeabilization, blocking, antibody incubation, washing and clearing, were executed using the automated EL406 liquid-handing system and Integra ViaFlo, ensuring consistency and reproducibility across experiments. Primary and secondary antibody information are described in Table S2.

### Tumoroid 3D 4i-based clearing wholemount staining

Fixed samples were equilibrated before staining. Permeabilization was performed by dispensing a permeabilization buffer (4% Triton X-100, 1% Tween-20 in PBS). Samples were incubated overnight at 4°C. Blocking was carried out using a buffer containing NH_4_Cl and Maleimide to minimize background staining. Blocking buffer (6% Donkey Serum, 1% BSA, 4% Triton X-100, 1% Tween-20, 100mM NH_4_Cl, 300mM Maleimide) was dispensed using the EL406 system and incubated for a total of 5–8 hours at room temperature in sequential incubation steps. Primary antibodies were diluted in blocking buffer without NH_4_Cl/Maleimide and incubated at 4°C. Following staining, samples were washed multiple times with blocking buffer without NH_4_Cl/Maleimide (1×PBS-based buffer). Optical clearing was performed using a sucrose clearing solution (glycerol, fructose, NAC) to enhance imaging depth. Samples were incubated in a clearing buffer at room temperature until transparency was achieved in 2–4 hours. For multiplexed imaging, antibodies were eluted using a glycine-urea-based stripping buffer (500mM Glycine, 3M Urea, 3M Guanidine chloride, 70mM TCEP in dH_2_O) before initiating the next round of staining. The entire workflow was automated using the EL406 liquid-handling system to ensure consistency and reproducibility. The original 2D cell culture-based 4i method (Gut et al.) was adapted and optimized for this 3D whole-mount clearing-based 4i method.

### Tumoroid time course assay

Samples from the co-cultured tumoroid models were collected at multiple time points (day 4, 7, 10 and 14) and 12 distinct antibody staining sets (Table S2) to evaluate temporal changes in cellular composition and morphology. At each time point, samples were fixed with 4% PFA at room temperature for 45 minutes and subsequently washed thoroughly with PBS to remove residual PFA. Fixed samples underwent staining and imaging following the established tumoroid staining and image acquisition workflow. This included permeabilization, blocking, antibody incubation, washing, and optical clearing using a fructose based clearing buffer. Imaging was performed post clearing using high-resolution microscopy to access structural and molecular changes over time. All steps, including cell culture, fixation, staining were performed using the automated EL406, Integra ViaFlo system, ensuring reproducibility and high throughput analysis across different time points.

### Tumoroid pilot screen and treatment condition optimization

Inhibitor perturbation screen was performed at multiple time points, with treatment initiation at day 4, 7, and 10 and the final endpoint at day 14. Tumoroids were treated with a panel of small molecule inhibitors at various concentrations (1, 5 and 10 µM), as detailed in Table S3. Fresh inhibitors were added daily during medium changes to maintain consistent drug exposure. The automated liquid-handling system facilitated precise dosing and high throughput screening, ensuring reproducibility across experimental conditions. The impact of inhibitor treatment was evaluated at the endpoint using 4i-based 3D clearing wholemount imaging for morphological analysis, allowing a comprehensive assessment of cellular responses to treatment.

### Tumoroid compound screen

Compound screening assay was conducted using a library of 650 small-molecule compounds (Table S4), each stored as a 2 mM DMSO stock in 384-well plates. Co-cultured tumoroids were prepared in 384-well plates as described above. Compound treatment began on day 4 and continued until day 14. Each plate included a DMSO control to account for solvent effects. Using the Integra ViaFlo system, 40 µL of a 10 µM intermediate compound dilution was added to the tumoroid medium, achieving a final compound concentration of 5 µM. Fresh compounds were replenished daily alongside the medium change to ensure sustained exposure. The automated liquid-handling system facilitated precise dosing and high-throughput screening, ensuring reproducibility across experimental conditions. The impact of compound treatment was evaluated at the endpoint using 4i-based 3D clearing whole-mount imaging for morphological analysis, enabling a comprehensive assessment of cellular responses to treatment.

To guide compound selection, differential gene expression (DE) analysis was performed on scRNA-seq data from the CAFs co-cultured tumoroids which was profiled over time (day7-day14). Additionally, benchmarking used the human PDAC biopsy and liver metastasis scRNA-seq datasets (Peng et al., a; Raghavan et al.) to confirm that tumoroid progression recapitulated in vivo trajectories. Differentially expressed genes (DEGs) had adjusted p-value < 0.05, AUC score > 0.6, log2 fold change > 0.15, were detected in at least 30% of the tested group and detection rate in the tested group was at least 30% compared to the background. The statistical test used was WilcoxSumRank. This analysis yielded the initial gene list for compound prioritization. Pathway enrichment (KEGG) of the DEG list identified functional modules, and compounds were included if their annotated targets mapped directly to DEGs or to enriched pathways. Redundancy was reduced to maximize unique target coverage, yielding the initial compound library.

Combinatorial therapeutic assay with small-molecule inhibitor and TCB in tumoroid tri-cultures. Co-cultures of tumoroids with cancer cells, CAFs, and TILs were established as described above. On day 11 of culture, treatments were initiated with a combination of TROP2–CD3 bispecific antibody (TROP2-TCB, 10 µg/ml) and the MEK inhibitor Trametinib (5 µM). A matched non-targeting control TCB (containing a CD3-binding arm and a non-specific DP47 arm) was used in parallel under identical conditions. An automated liquid-handling system (specify instrument/model) was used to deliver precise dosing and facilitate high-throughput application, ensuring reproducibility across experimental replicates. The treatment regimen was maintained until day 14, at which point cultures were analyzed. Therapeutic impact was evaluated using iterative indirect immunofluorescence imaging 4i-based 3D clearing wholemount imaging to enable high-resolution morphological and phenotypic assessment of cellular responses within the tri-culture microenvironment.

### Explant sliced tissue culture

Tru-cut biopsies from pancreatic tumors were received on ice within 24 hr from surgery. All tissue processing was carried out on ice to maintain tissue viability. Biopsies were washed in a cold wash medium (advanced DMEM/F12 + 1:250 Primocin + 2% PenStrep) and kept submerged in this medium until the moment of further processing. Biopsies were processed in one of two ways: For biopsies with good length (>5mm), tissue cores were arranged within a pre-wetted tissue slicer matrix (Zivic Instruments, HSMS005-2) such that their longer dimension was perpendicular to the direction of cutting within the matrix. Razor blades were snapped in two, pre-wetted in ice-cold wash medium and inserted, blade-side first, into the slicer matrix sequentially, such that they would rest on the tissue cores. Once all blades were inserted, they were collectively pushed downwards in one clean motion, thereby cutting each tissue core into several (4–8) 0.5 mm-thick slices. Tissue slices were then collected from each blade individually using tweezers and maintained in a wash buffer until embedding on transwells. For biopsies with poor tissue integrity (<0.5 mm length), slices were generated using a Krumdieck tissue slicer (Alabama R&D, MD6000), according to the manufacturer’s protocol. In short, multiple tissue cores were embedded in agarose solution (2% w/v) within refrigerated molds (Alabama R&D, MD2200). The resulting tissue-agarose composite cores were inserted into the Krumdieck sample holder, submerged in ice-cold PBS and sliced to 0.5 mm thick slices. Towards this, the arm and blade vibration speed were set to medium. Collected slices were returned to the wash buffer, freed from the surrounding agarose using tweezers and maintained cold until further embedding on transwells. The sectioned tissue slides were cultured in the same medium formulation used for the tri-culture of tumoroids with cancer cells, CAF and TILs. TROP2-TCB (10 µg/ml) and Trametinib (5 µM) were treated from day 0 of culture, with daily medium replacement containing the respective treatments, and maintained continuously until day 4.

### FFPE embedding of co-cultured tumoroid

Samples were washed three times with 1×DPBS before fixation with 4% paraformaldehyde (PFA) in the 96-well Clear TC-treated plate. Following 45 minutes of fixation at room temperature, the wells were washed three more times before complete aspiration of 1×DPBS. Preliquefied HistoGel (ThermoScientific) was distributed into biopsy cassettes then samples were corrected from the 96-well Clear TC-treated plate then embedded into HistoGel before completely polymerized. Samples were dehydrated overnight using a Vacuum filter processor (Sakura, TissueTek VIP5). The following day, samples were embedded in liquid paraffin. FPE blocks were generally sectioned at a thickness of 3.5 µm and mounted on Superfrost Plus Adhesion microscope slides (Epredia). Slides were incubated overnight at 37°C in a slide oven.

### FFPE-based H&E staining

Slides were deparaffinized, rehydrated to distilled water, and stained with hematoxylin (Sigma-Aldrich) for 5 minutes, followed by rinsing under running water for 5 minutes. Sections were then differentiated in 0.3% acid alcohol for 20 seconds and washed again under running water for 5 minutes. Counterstaining was performed with eosin (Thermo Fisher) for 2 minutes, after which slides were dehydrated, cleared, and mounted.

### FFPE-based immunohistochemistry staining

FFPE sections on dried slides were stained using a Ventana Discovery Ultra automated stainer (Roche Tissue Diagnostics). Antibodies are listed with Table S2. Slides were baked at 60°C for 8 minutes and deparaffinized in three cycles of Discovery Wash at 69°C for 8 minutes each. Heat-induced epitope retrieval was performed with Discovery CC1 for 40 minutes at 92°C, followed by incubation with Inhibitor CM (ChromoMap DAB kit, Roche) for 4 minutes at room temperature. Primary antibodies were applied at the indicated dilutions and incubated for 60 minutes at 37°C, after which HRP-conjugated secondary antibody (multimer HRP) was added for 16 minutes at room temperature. Signal detection was performed with DAB for 8 minutes. Nuclei were counterstained with hematoxylin for 8 minutes and bluing reagent for 4 minutes. Finally, slides were cleared in xylene and mounted.

### Brightfield imaging

H&E and IHC stained slides were imaged using a Hamamatsu NanoZoomer S360 whole-slide scanner at 20× magnification, yielding a pixel size of 0.23 µm per pixel for all brightfield images.

### FFPE-based multiplex immunofluorescence (mIF) staining

mIF staining of FFPE slides was performed on a Ventana Discovery Ultra automated tissue stainer (Roche Tissue Diagnostics). Slides were baked at 60°C for 8 minutes and then heated to 69°C for 8 minutes for deparaffinization; this cycle was repeated twice. Heat-induced antigen retrieval was carried out with Tris–EDTA buffer (pH 7.8; Ventana) at 92°C for 32 minutes. Each blocking step with Discovery Inhibitor (Ventana; 16 minutes) was followed by neutralization. Primary antibodies were diluted in Discovery Ab diluent (Ventana) and detected with species-specific HRP-conjugated secondary antibodies (OmniMap Ventana; Table S2), after which the appropriate Opal dye (Akoya Biosciences) was applied. Following each cycle of primary, secondary and Opal dye application, residual antibodies and HRP were removed through antibody neutralization and HRP denaturation before repeating the staining cycle with a new blocking step. Nuclei were counterstained with DAPI (Roche).

### FFPE-based mIF imaging

Stained slides were digitized using multispectral imaging on a Vectra Polaris system (PerkinElmer) with MOTiF technology at 20× magnification, capturing all seven fluorophores (Opal 480, 520, 570, 620, 690, 780 and DAPI). Slides were scanned in batches under identical imaging conditions to ensure comparability for downstream HALO AI (Indica Labs) analysis. Channel unmixing and image tiling were performed using PhenoChart (v1.0.12) and inForm (v2.4), and tiles were fused in HALO (v3.2.1851.328).

### Whole-mount immunofluorescence imaging, processing and analysis Image acquisition

In all samples, high-throughput imaging was done with an automated spinning disk microscope from Yokogawa (CellVoyager 8000), with an enhanced CSU-W1 spinning disk (Microlens-enhanced dual Nipkow disk confocal scanner), a 20x (NA = 1.0) Olympus objective, and a Neo sCMOS camera (Andor, 2000×2000 pixels). For imaging, an intelligent imaging approach was used (Search First module of Wako Software Suite (Fujifilm Wako Automation)). In brief, each well of muti-well plates was acquired as a single field of view using a 4x air objective (Olympus, NA = 0.45) in order to cover the complete well area. Generated images were used to segment objects on the fly with a custom ImageJ macro, outputting coordinates of individual organoid positions. These coordinates were used to generate a map of locations for high resolution re-imaging (20x water immersion, NA = 1.0). Four fields of view were used at 20x magnification to cover each object. Tiles were acquired with 10% overlap to facilitate stitching. For each site, z-planes spanning a range up to 600 µm were acquired. 10 µm z-step was used in all experiments unless indicated otherwise.

### Image preprocessing and background correction

Illumination correction was performed by computing flatfield images using the BaSiC method (Basic and Stable Illumination Correction) (Peng et al., b). For each plate, flatfields were estimated for all individual channels by sampling from all wells in batches of 250 images per channel and averaging the resulting flatfield images. Tile stitching utilized a 10% overlap between adjacent image tiles with linear blending in the overlap regions to create seamless transitions between fields of view. The stitching process accommodated configurations of 3×3 tiles per well. For background correction BaSiC was applied in additive or multiplicative correction mode depending on channel characteristics (smoothness_flatfield=1, max_reweight_iterations=1000). All processing steps were parallelized using Dask (Rock- lin) distributed computing on HPC infrastructure. Quality control metrics assessed signal consistency across wells by computing foreground-to-background ratios by Otsu thresholding (threshold=0.005) and channel-specific intensity statistics (mean, standard deviation, median and median absolute deviation).

### Message-passing features for spatial neighborhood analysis

To characterize spatial neighborhood patterns in multiplexed image data, a message-passing framework was developed that integrates neighborhood information based on segmented nuclei features and image information from tissue subregions between nuclei. This approach leverages graph representations of cellular proximity relationships to capture complex microenvironmental interactions. Similar to Kim et al. (b), a spatial interaction graph was constructed by connecting cells within a 100 pixel radius, establishing edges based on physical distance constraints. For each marker, cell-specific feature vectors were calculated by aggregating signals from directly connected neighbors through a normalized adjacency matrix multiplication, effectively passing information through the cellular network. This message-passing implementation extends traditional graph analysis by incorporating both self-connections and neighbor-weighted signals derived from oriented image subregions located along edges of the spatial graph. Given a binary adjacency matrix *A* representing spatial cellular connectivity and a feature matrix *X* containing marker intensities, a weighted adjacency matrix *A*_*weight*_ = *D*_*inv*_ × *A* was derived, where *D*_*inv*_ is the inverse degree matrix ensuring each cell’s neighborhood contribution is normalized. The message-passed features were then computed as *M* = *A*_*weight*_ × *X*, creating a contextualized representation for each cell that incorporates its spatial neighborhood characteristics. For edge-specific feature aggregation, oriented sampling windows were placed in the actual image space along the physical paths connecting adjacent nuclei, with each window having a width of 10 pixels and following the angle of the connection. This generated two complementary feature representations: cell-intrinsic message-passed features (neighborhood-integrated) and features that incorporate signals between cells (directionally-weighted) which were weighted equally to derive the final feature matrix.

### Nuclei segmentation and feature extraction

Nuclear segmentation employed Cellpose3 (Stringer and Pachitariu) with model_type=cyto3, diameter=25, flow_-threshold=0.5, cellprob_threshold=0. For DAPI channel restoration, Cellpose’s denoise model was utilized. Label merging across z-stacks (2.5D) utilized Cellpose’s stitch3D function to establish consistent 3D objects by resolving overlapping regions. Features were extracted using skimage’s regionprops_table, measuring morphological properties (area, eccentricity, major/minor axis length) and intensity distributions (mean, standard deviation, median and median absolute deviation) for each channel. Additionally, the message-passing approach described above was applied with 100 pixel radius sampling to characterize spatial cellular relationships. Cell feature and respective message-passed data was organized in a data format with position coordinates, intensity measurements, and morphological properties stored as observations, enabling compatibility with analytical frameworks like Scanpy (Wolf et al.) and Muon (Bredikhin et al.). Ridge regression was applied to correct for photobleaching effects (Polański et al.) along the z-axis, utilizing BaSiC-derived transformation matrices for consistent intensity normalization across imaging rounds.

### Image analysis tumoroid tri-cultures

Imaris (Oxford Instruments, v10.1.1) was used for visualization and quantitative analysis of 3D immunofluorescence images. Channel arithmetic was applied to reduce bleed-through between fluorophores. Immune cells were segmented using the Spots function. Fibroblasts (Collagen I+) were quantified by nuclei overlap within Collagen I positive regions. Tumor growth was assessed by measuring the total 3D tumoroid volume. Morphological parameters including volume, sphericity, and position were quantified, and visualized using GraphPad Prism10.

### Phenocoder

#### Model architecture

Phenocoder implements a conditional variational autoencoder (cVAE) framework for learning latent representations of multiplexed whole-mount imaging data. The encoder transforms 128×128×n dimensional input image patches (timecourse data: *n*=1, screening data: *n*=4) into a 32-dimensional latent space, conditioned on experimental variables to control for technical variation.

The encoder consists of a series of convolutional layers with filter sizes (64, 128, 256, 512), 3×3 kernels, and stride 2, each followed by ReLU activation. The flattened features are concatenated with one-hot encoded condition variables *c* (plate ID and z-stack position) before passing through a dense layer (128 neurons) with 25% dropout regularization.

Two parallel dense layers output the latent distribution parameters *µ*(*x, c*) and log *σ*^2^(*x, c*), parameterizing the approximate posterior *q*_*ϕ*_(*z* | *x, c*) = 𝒩 (*µ, σ*^2^*I*). Latent vectors *z* are sampled using the reparameterization trick (Kingma and Welling):

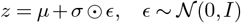

The decoder architecture mirrors the encoder, starting with a dense layer that concatenates the latent vector *z* with condition variables *c*, then reshapes to 8×8×64 dimensions. This is followed by four transposed convolutional layers (3×3 kernel, stride 2, ReLU activation) with matching filter sizes in reverse order. A final transposed convolution with sigmoid activation produces the reconstructed image patches 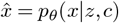. The total loss function combines reconstruction error and Kullback–Leibler (KL) divergence:

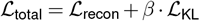

where the reconstruction loss uses binary cross-entropy (BCE) summed over spatial dimensions:

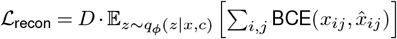

with *D* = *n* channels of the respective datasets. The KL divergence term regularizes the latent space:

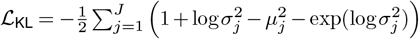

where *J* is the latent dimension. The weighting factor *β*=0.01 prevents posterior collapse while prioritizing reconstruction fidelity.

The model was implemented in Python 3.10.14 using TensorFlow as the backend and Keras for model construction. For implementation details and code, see: https://github.com/devsystemslab/pdac_tumoroid

### Dataset generation

Multiplexed whole-mount images were processed to extract 128×128 pixel patches centered on detected nuclei across all imaging channels. The extraction process targeted image patches within the boundaries of the images, ensuring complete patch coverage. Signal intensity was normalized using percentile-based scaling (1st–99th percentiles) to standardize ranges across different markers. For model training and evaluation, the dataset was split by assigning complete tissue wells to either training (80%) or validation (20%) sets, ensuring sample separation to prevent data leakage while maintaining representation of biological heterogeneity.

### Training

The model was trained using Adam optimizer with an initial learning rate of 0.001 on 80% of the data. Training proceeded for up to 50 epochs with three key callbacks implemented: early stopping with patience of 5 epochs monitoring validation loss, learning rate reduction (factor 0.2, patience 3 epochs) when performance plateaued, and TensorBoard logging for performance tracking. The best-performing model weights were restored based on validation loss. After training, model performance was evaluated by observed patch reconstruction errors and were assessed for batch effect correction and biological variability.

### Unsupervised clustering

After model training, all nuclei-centered patches were encoded into the latent space using the learned encoder network. For nuclei appearing across multiple z-slices, only the middle slice was selected to ensure consistent representation. Unsupervised clustering using the Leiden algorithm (resolution=0.5) was performed on a representative subset of one million encoded cells through a nearest-neighbor graph (k=15). Cluster assignments were then propagated to the complete dataset using PyNNDescent (Dong et al.) for efficient nearest-neighbor search.

### Benchmarking

To quantitatively assess the preservation of spatial cellular organization in latent representations, complementary metrics that evaluate distinct aspects of spatial information retention across four embedding approaches were implemented for the time course data and the pilot screen data: (1) nuclei intensity features derived directly from immunofluorescence channels combined with morphological measurements, (2) message-passed nuclei features that incorporate local neighborhood context, (3) Phenocoder latent embeddings learned through the conditional variational autoencoder, and (4) message-passed Phenocoder embeddings that combine deep representation learning with spatial context integration. These methods were compared using the following metrics:

1. Graph Connectivity Similarity (GCS): Quantifies how accurately the latent nearest neighbor graph preserves edges and non-edges from the spatial nearest neighbor graph. Connectivity matrices were compared using normalized Frobenius norm of element-wise differences between adjacency matrices. Values range from 0 (no preservation) to 1 (perfect preservation), providing a global measure of structural similarity between latent and spatial representations.
2. Maximum Leiden Adjusted Mutual Information (MLAMI): Evaluates global spatial organization preservation through clustering consistency. Leiden clustering was computed at multiple resolutions (0.1–1.0) for both spatial and latent representations, then calculated the Adjusted Mutual Information (AMI) between all clustering resolution pairs. The maximum AMI across all resolution combinations served as our measure of global structure preservation.
3. Cell Type Local Inverse Simpson’s Index Similarity (CLISIS): Assesses preservation of local neighborhood heterogeneity between spatial and latent representations. Cell Type Local Inverse Simpson’s Index was computed on both nearest neighbor graphs, took their ratio, and applied appropriate normalization to obtain a metric ranging from 0 to 1, with higher values indicating better preservation of local neighborhood structure.
4. Niche Average Silhouette Width (NASW): Measures the separability and compactness of clusters in the latent space. Leiden clustering was computed at multiple resolutions, calculated the silhouette width for each clustering, and averaged them to quantify the distinctness of identified cell niches in the latent space.
5. Adjusted Rand Index (ARI): To assess consistency of unsupervised cell-type identification,the Adjusted Rand Index between clustering results from different embedding methods was computed. For each embedding approach, Leiden clustering was performed with identical parameters (resolution=0.5, n_neighbors=15). ARI values range from −1 to 1, with 1 indicating perfect agreement between clusterings, 0 indicating random chance agreement, and negative values indicating agreement worse than random chance.

Values for each metric were computed on a per-organoid basis and aggregated across the entire dataset. These benchmarks were implemented using a modified framework derived from NicheCompass (Birk et al.).

### Phenotypic organoid embeddings

Organoid-level embeddings based on microenvironmental latent representation were generated to quantify drug-induced phenotypic changes. Spatial graphs connecting nuclei within 100 pixels were constructed based on physical coordinates, enabling message passing across neighboring cells to generate context-aware representations. Cell neighborhood clusters were annotated by integrating latent embedding features with immunofluorescence intensities and morphological measurements (eccentricity, area, major and minor axis length). Organoid-level representations were derived from both original and message-passed embeddings through Leiden clustering. Spatial statistics were calculated using Squidpy (Palla et al.), including spatial autocorrelation (Moran’s I) for marker expression, interaction scores between cell clusters, and neighborhood enrichment analyses. All summary statistics dependent on spatial graph representations were calculated for 25, 50, 100 and 150 pixel maximal edge lengths. Tumoroid volume, surface area and density were approximated by constructing a spatial graph with a maximum radius of 100 pixels, pruning the graph to include only nodes with a degree greater than 5, and calculating the convex hull. For the pilot screen data, ductal structures were identified as connected components of cancer cell clusters, with volumes calculated using convex hulls on pruned spatial graphs. Additional feature extraction included distance metrics of segmented ducts to the center of mass of all nuclei centroids, LAMC2 and KI67 activity scores, fibroblast-epithelial interaction metrics, per-organoid duct count, cell density in ducts, and total duct area. These multidimensional features collectively formed the organoid phenospace which after dimensionality reduction by PCA (PCs=32) were used for perturbation analysis of the screening data and spatiotemporal analysis of the time course data.

### Dataset specific analysis

#### KNN imputation and phenotypic trajectory in timecourse dataset

To impute all markers (Table S2) across all registered cells in the dataset, cells were grouped by stain marker and nearest-neighbor (NN) search (n=3) using PyNNDescent was performed on the PCA dimension-reduced space of the latent embedding, excluding self-indexes from NN. Marker intensity imputation was performed efficiently for each stain by computing the dot product between the adjacency matrix constructed from the NN search tree and the matrix of IF intensity values of cells in the marker group. Imputation was performed for nuclei and message-passed spatial microenvironment IF values. Imputation quality was assessed using 10-fold cross-validation with complete sample hold-out. Unsupervised Leiden clustering (res=0.5) was applied to all feature sets and whole-tumoroid embeddings were constructed as described above and a diffusion pseudotime trajectory (Haghverdi et al.) was computed. Phenotypic tumoroid features including volume, density, per cell type aggregated interaction terms, and spatial autocorrelation values (Moran’s I) at 150 pixel radius were visualized as locally weighted scatterplot smoothing (LOWESS) splines along the pseudotemporal-trajectory. Hierarchical clustering (Ward.D2) of Moran’s I values for Phenocoder clusters and imputed immunofluorescence intensities, and respective Spearman correlations was applied to reveal coordinated patterns of spatiotemporal organization dynamics during tumoroid maturation.

#### Effect size quantification in screening datasets

To quantify compound-induced phenotypic changes in tumoroid co-cultures, grouped Mahalanobis distances between treatment and DMSO control conditions in the PCA-reduced feature space was calculated. This approach accounts for covariance structure in the high-dimensional data, providing a statistically robust measure of phenotypic divergence. For each compound, we defined:

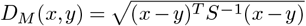

where *x* represents the centroid of the treatment group, *y* represents the centroid of the DMSO control group, and *S* is the pooled covariance matrix. This distance metric accounts for variable correlations and scales appropriately with feature dimensionality.

For the pilot screen, treatment effects were visualized as a heatmap using hierarchical clustering (Ward.D2) based on the determined Mahalanobis distances. To assess assay quality, linear discriminant analysis (LDA) was performed between all treatment conditions and DMSO controls, calculating *Z*^*′*^ factor values:

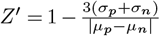

where *σ*_*p*_ and *σ*_*n*_ denote standard deviations, and *µ*_*p*_ and *µ*_*n*_ denote means of positive and negative controls, respectively.

For the tumoroid compound screen, this approach was extended to evaluate 650 compounds targeting diverse cellular mechanisms. The dataset was filtered to exclude wells with imaging artifacts. Compound effects were aggregated at gene target level by calculating weighted distances:

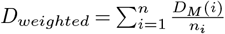

where *D*_*M*_ (*i*) represents the Mahalanobis distance for compound *i* and *n*_*i*_ is the number of compounds targeting the same gene. This approach normalized for gene targets with multiple selective inhibitors.

### Integrated analysis of phenotypic space

To connect phenotypic effects with primary tissue PDAC biology, the scRNA-seq atlas was integrated by examining target gene expression across cellular compartments in primary tumors. Compound targets were ranked based on phenotypic effect sizes, and pathway enrichment analysis was performed using a hypergeometric test. This enabled identification of biological processes most susceptible to perturbation in the cancer-fibroblast interactome.

To visualize the functional landscape of druggable targets in the tumor-stroma microenvironment, a STRING protein-protein interaction network (Szklarczyk et al.) of the top 100 gene targets ranked by weighted distance was constructed. STRING’s unsupervised clustering (DBSCAN) was applied to group targets into functional modules. Pathway annotations were overlaid onto the network using color coding, with node size mapped to phenotypic effect magnitude.

## Supplementary Information

**Extended Data Figure 1.**
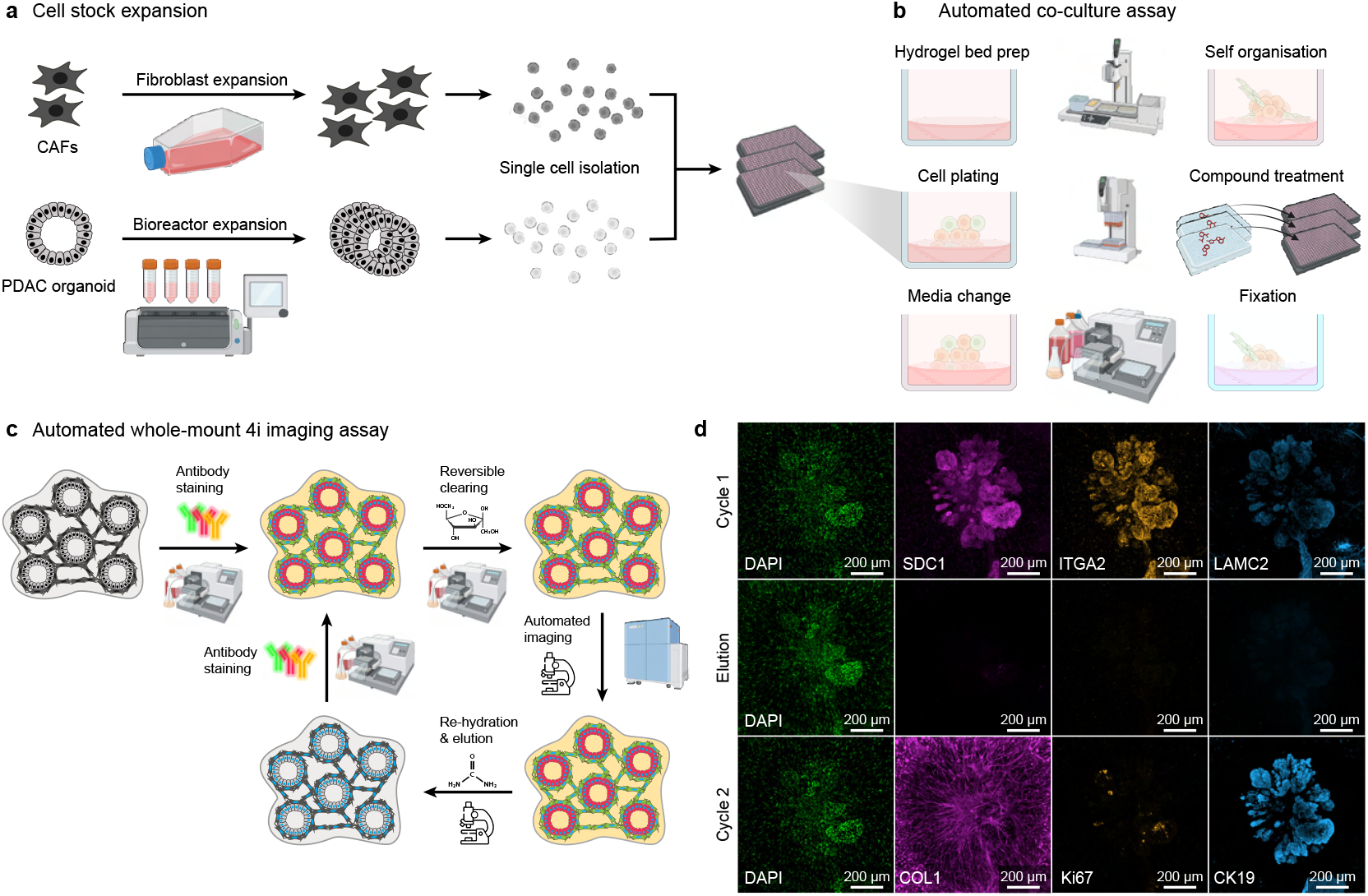
Tumoroid culture and multiplexed high-throughput imaging automation. a) Cell stock expansion of cancer-associated fibroblasts (CAFs) and pancreatic ductal adenocarcinoma (PDAC) organoids to establish PDAC organoids for downstream assays. b) In the automated co-culture assay PDAC organoids and CAFs are seeded as single cells on hydrogel beds prepared in multiwell plates. The automated workflow includes seeding of cells, media exchange, compound treatment, and fixation for endpoint imaging. c) Automated whole-mount 4i imaging assay for multiplexed molecular profiling of co-cultures. Fixed samples are first stained with antibodies specific to desired molecular targets, followed by reversible clearing with glycerol-fructose buffer and automated imaging to capture fluorescence signals. Imaging is followed by elution of bound antibodies, enabling iterative rounds of staining, clearing, and re-imaging to detect multiple markers.

**Extended Data Figure 2.**
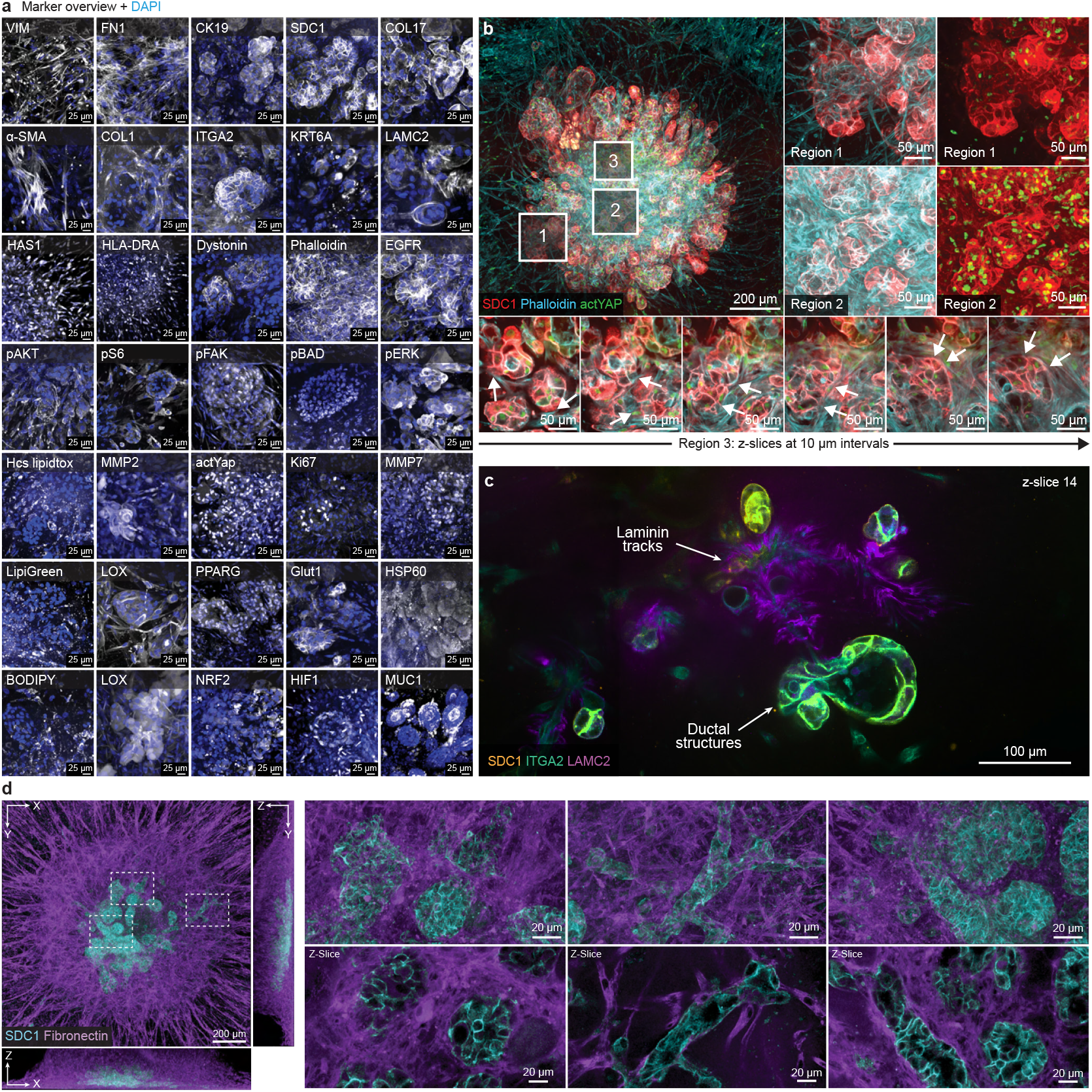
Extended phenotyping of dynamics and structural features in 3D human fibrotic pancreatic tumoroids. a) Immunofluorescence profiling of day 14 tumoroids showing expression patterns of 35 different markers (white) and DAPI stain (blue), including structural proteins, cell-type markers, adhesion molecules, signaling pathway components, metabolic markers, and functional indicators. b) Fibrotic tumoroid stained with SDC1 (red), Phalloidin (cyan), and actYAP (green). Three regions of interest (1-3) are magnified to show detailed tissue architecture at the duct-stroma interface. Region 3 is visualized as sequential z-slices at 10 µm intervals, arrows indicating interaction interfaces between cancer and ECM. c) Tumoroid (z-slice 14) stained with SDC1 (orange), ITGA2 (green), and LAMC2 (magenta). d) 3D (X-Y-Z) visualization of tumoroid stained with SDC1 (cyan) and fibronectin (magenta). Orthogonal projections and high-resolution z-slices illustrate the spatial organization of epithelial and stromal compartments.

**Extended Data Figure 3.**
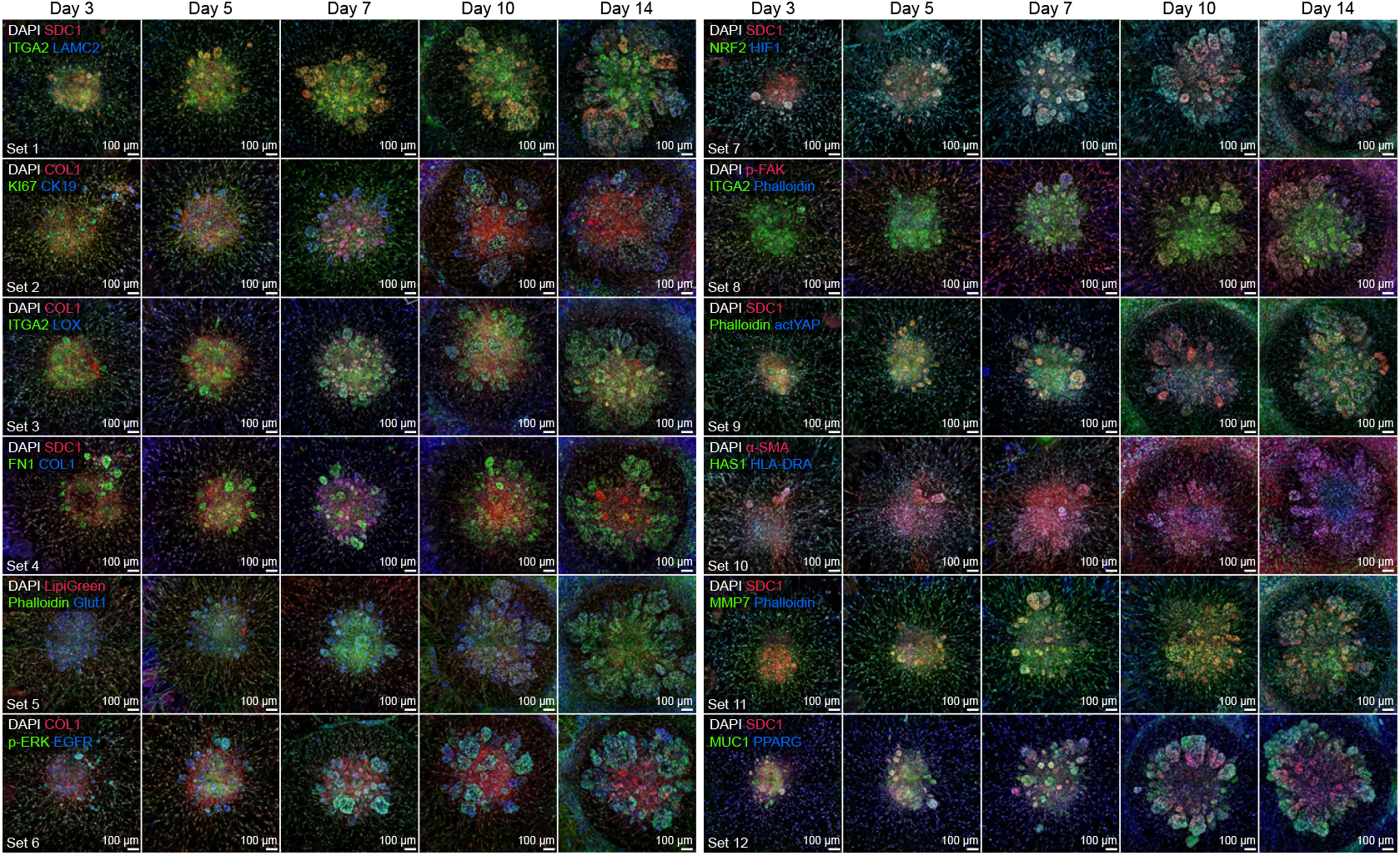
Temporal dynamics of tumoroid development across multiplexed immunofluorescence staining panels. Montage overview showing temporal progression of tumoroid development from day 3 to day 14 across 12 multiplexed staining sets. Each row displays a specific antibody combination (Table S1) in combination with DAPI, revealing dynamic changes in cancer cell markers, stromal components, matrix proteins, and signaling pathways during tumoroid maturation. Representative images demonstrate the progression of distinct architectural features including ductal structures, fibrotic cores, and cancer-stromal interfaces.

**Extended Data Figure 4.**
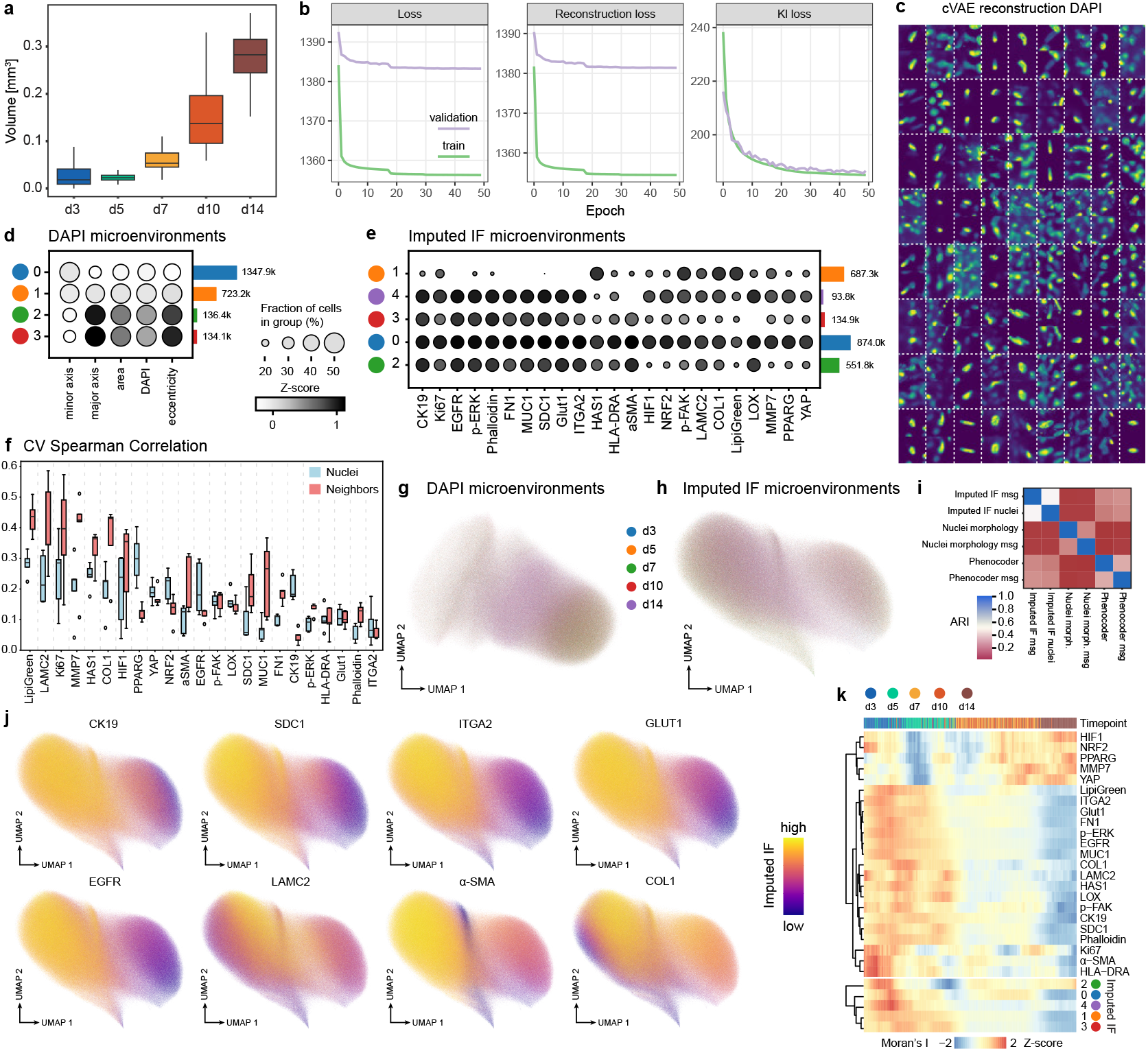
Phenocoder analysis of timecourse data. a) Boxplots show tumoroid volume [mm^3^] progression at day 3, 5, 7, 10, and 14. b) Training history plots showing the convergence of loss functions during cVAE training for 50 epochs. Metrics include total loss, reconstruction loss, and KL divergence loss for training (green) and validation (purple) set. c) Visual comparison of input (left) and reconstructed (right) DAPI image patches (128×128 pixels). d,e) Dotplots quantifying feature distributions across Phenocoder clusters of microenvironments based on DAPI or imputed immunofluorescence. (d) Shows the relative expression of DAPI and measurements of morphological features (eccentricity, minor/major axis, area), and (e) marker expression across all imputed markers. Circle size represents the fraction of cells in each group, with the number of cells per cluster indicated as barplots. f) Barplots show Pearson’s correlation coefficient for imputed marker distributions (nuclei and spatial neighbors) across all marker sets. g,h) UMAP embeddings for DAPI and imputed immunofluorescence microenvironments colored by timepoint. f) Barplots show Spearman’s correlation coefficient for imputed marker distributions (nuclei and spatial neighbors) across all marker sets from cross validation (CV) with sample hold outs. i) Heatmap shows Adjusted Rand Index (ARI) scores between clustering results from different embedding methods. These include DAPI based Phenocoder, nuclei based DAPI and morphology, and imputed IF embeddings and their respective embeddings after applying spatial message-passing. j) UMAP feature plots for selected epithelial and stromal imputed protein markers. k) Heatmap visualizing Z-scores of spatial autocorrelation values (Moran’s I) for imputed immunofluorescence and respective Phenocoder clusters along the pseudotemporal trajectory.

**Extended Data Figure 5.**
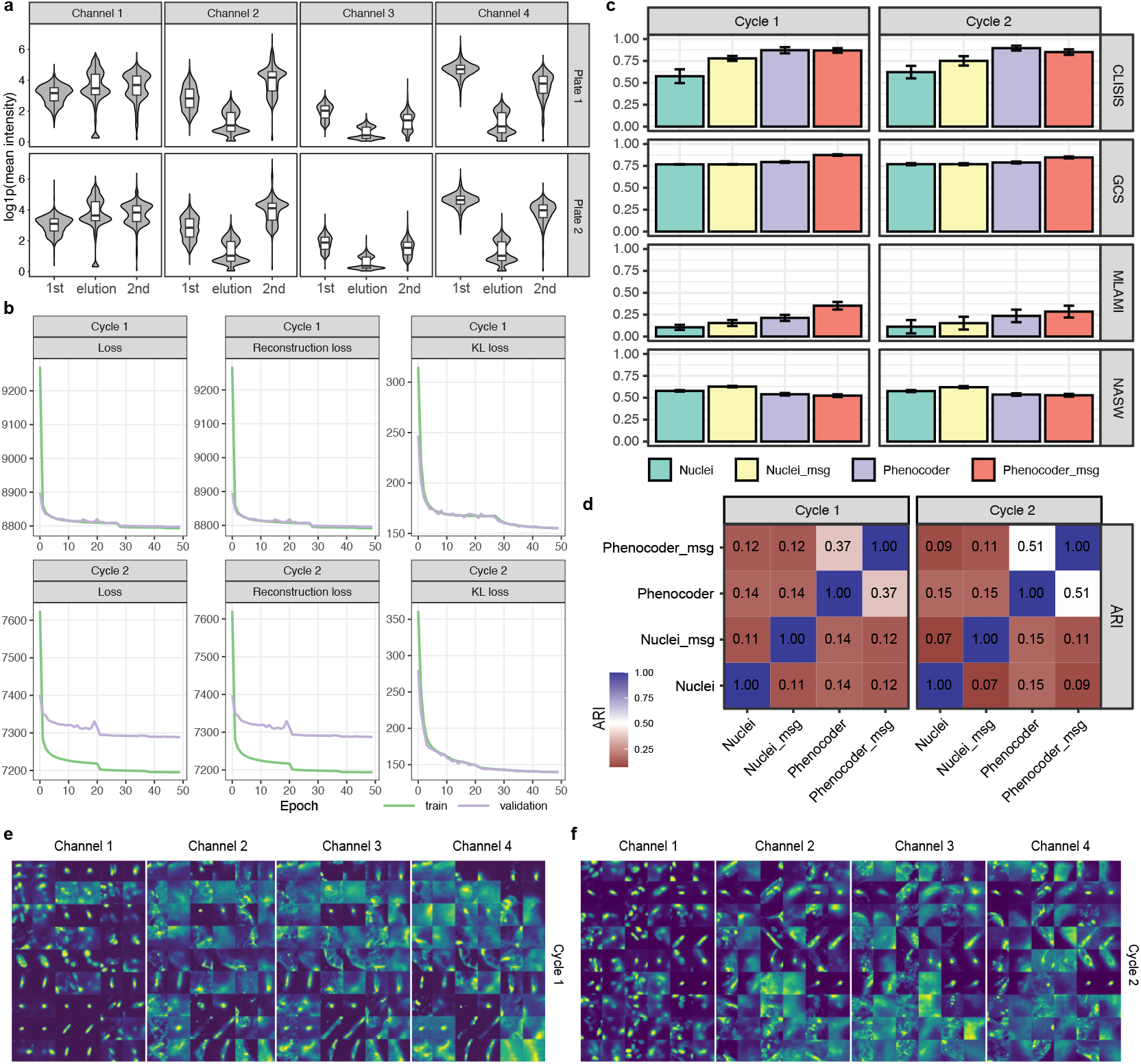
Quality assessment of pilot screen dataset. a) Violin plots with overlaid boxplots showing log1p-transformed mean intensity distributions across imaging channels (1-4) for two plates. Comparision across the first imaging, post-elution, and second imaging cycle, demonstrates signal recovery and consistency across imaging cycles. b) Training history plots showing the convergence of loss functions during cVAE training across 50 epochs. Metrics include total loss, reconstruction loss, and KL divergence loss. Green lines indicate training set performance while purple lines show validation set performance. c) Quantitative comparison of spatial conservation metrics between four different cluster label sets derived from raw nuclei features, message-passed nuclei features, Phenocoder embeddings, and message-passed Phenocoder embeddings. d) Heatmap showing Adjusted Rand Index (ARI) scores between clustering results from different embedding methods. e,f) Visual comparison of input (left) and reconstructed (right) image patches (128×128 pixels) from the cVAE across all four channels for cycle 1 (d) and cycle 2 (e).

**Extended Data Figure 6.**
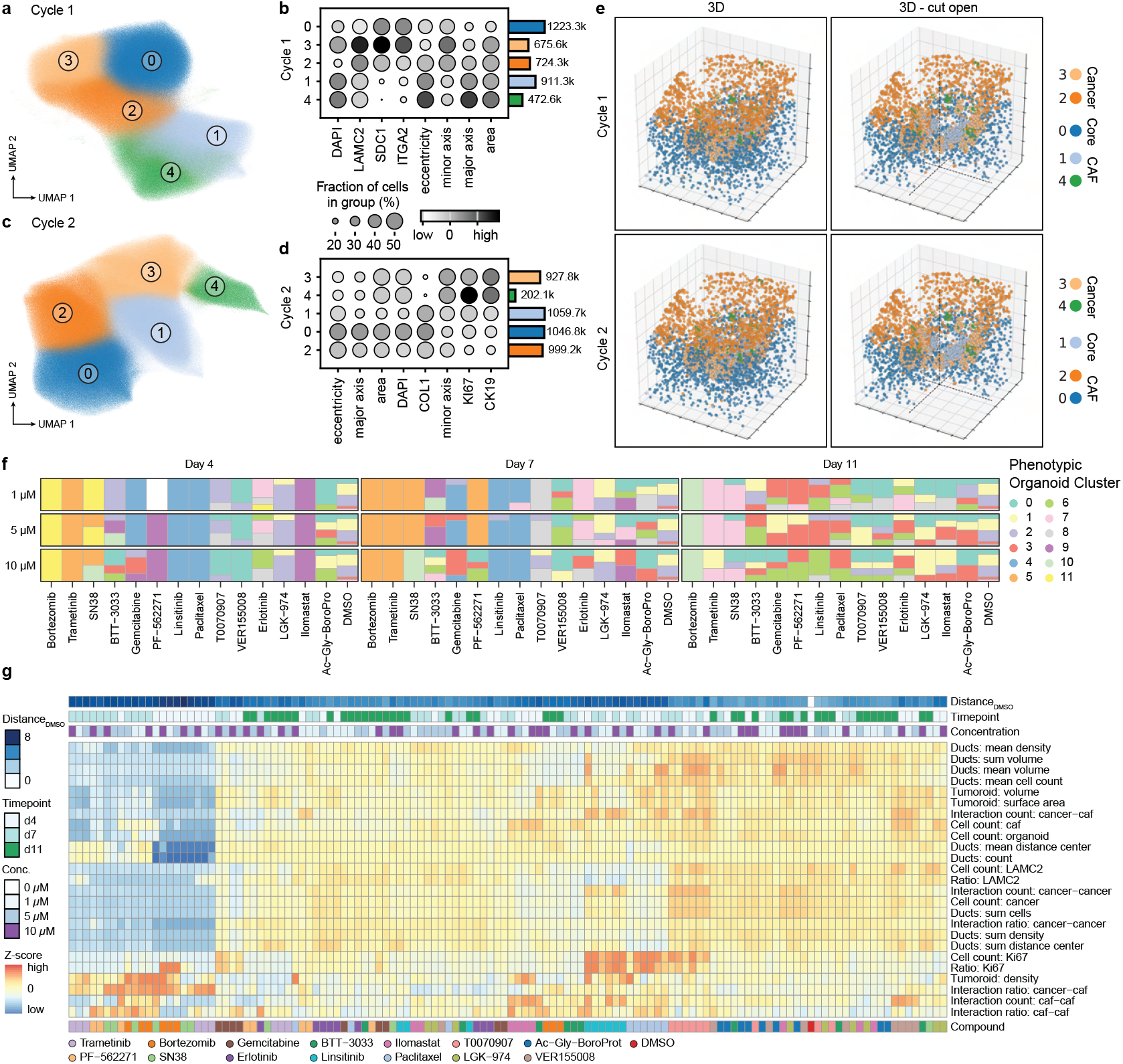
Extended Phenocoder analysis for the pilot screen. a,c) UMAPs of latent representations of nuclei-centered image patches (z) encoded by Phenocoder for imaging cycle 1 (a) and cycle 2 (c), both showing 5 distinct cellular phenotypic clusters (0-4) b,d) Dotplots quantifying feature distribution across Phenocoder clusters for cycle 1 (b) and cycle 2 (d). Cycle 1 shows the relative expression of DAPI, LAMC2, SDC1 and ITGA2, while cycle 2 shows DAPI, COL1, Ki67 and CK19. Both include measurements of morphological features (eccentricity, minor/major axis, area). Circle size represents the fraction of cells in each group (%), with number of cells per cluster indicated (right). e) 3D spatial visualization of encoded nuclei colored by Phenocoder cluster identity across both imaging cycles, shown in closed (left) and cut-open (right) perspectives to reveal internal organoid architecture. f) Stacked barplots showing distribution of phenotypic organoid clusters (0-10) across all compound treatments, organized by concentration (1, 5, 10 µM) and timepoint (Day 4, 7, 11). g) Heatmap visualizing a selection of aggregated features (Z-score) of the latent phenotypic organoid embedding organized. Aggregated features include: structural metrics (ductal density, volume, distance); cellular quantification (cell counts by type); marker expression (LAMC2 and Ki67); and interaction metrics (cancer-CAF, cancer-cancer, CAF-CAF spatial relationships). Additional annotation with distance to DMSO controls color-coded annotation bars indicate compound identity, timepoint, and concentration.

**Extended Data Figure 7.**
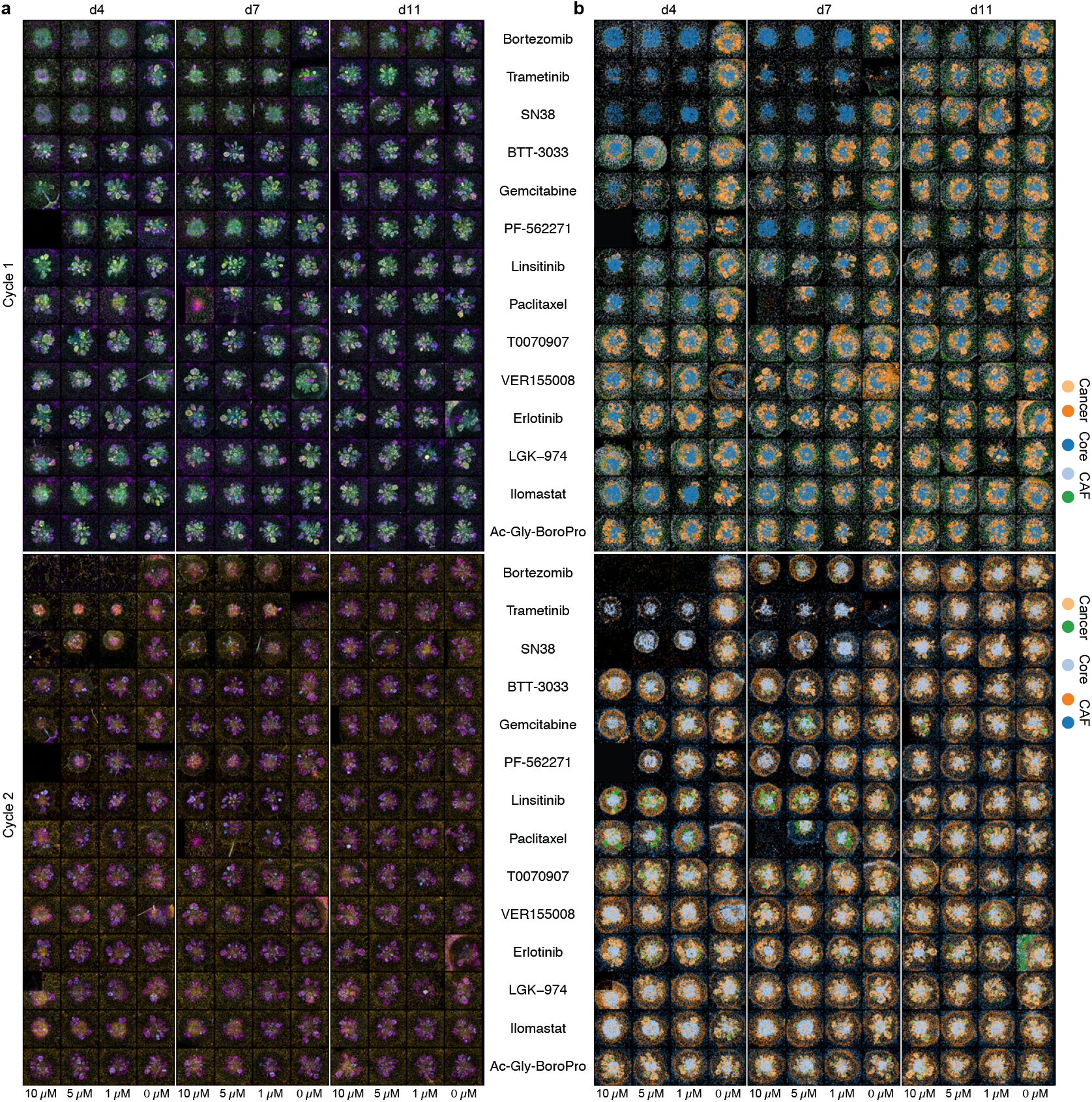
Overview montage of whole-mount immunofluorescence and Phenocoder clusters for all treatment conditions. a) Max-projections of whole-mount immunofluorescence images of tumoroids treated with 14 different drugs at varying concentrations (10 µM, 5 µM, 1 µM, 0 µM) over three timepoints (days 4, 7, 11) across two experimental cycles. b) Corresponding Phenocoder cluster analysis visualizing drug-induced phenotypic changes in the tumoroid co-cultures.

**Extended Data Figure 8.**
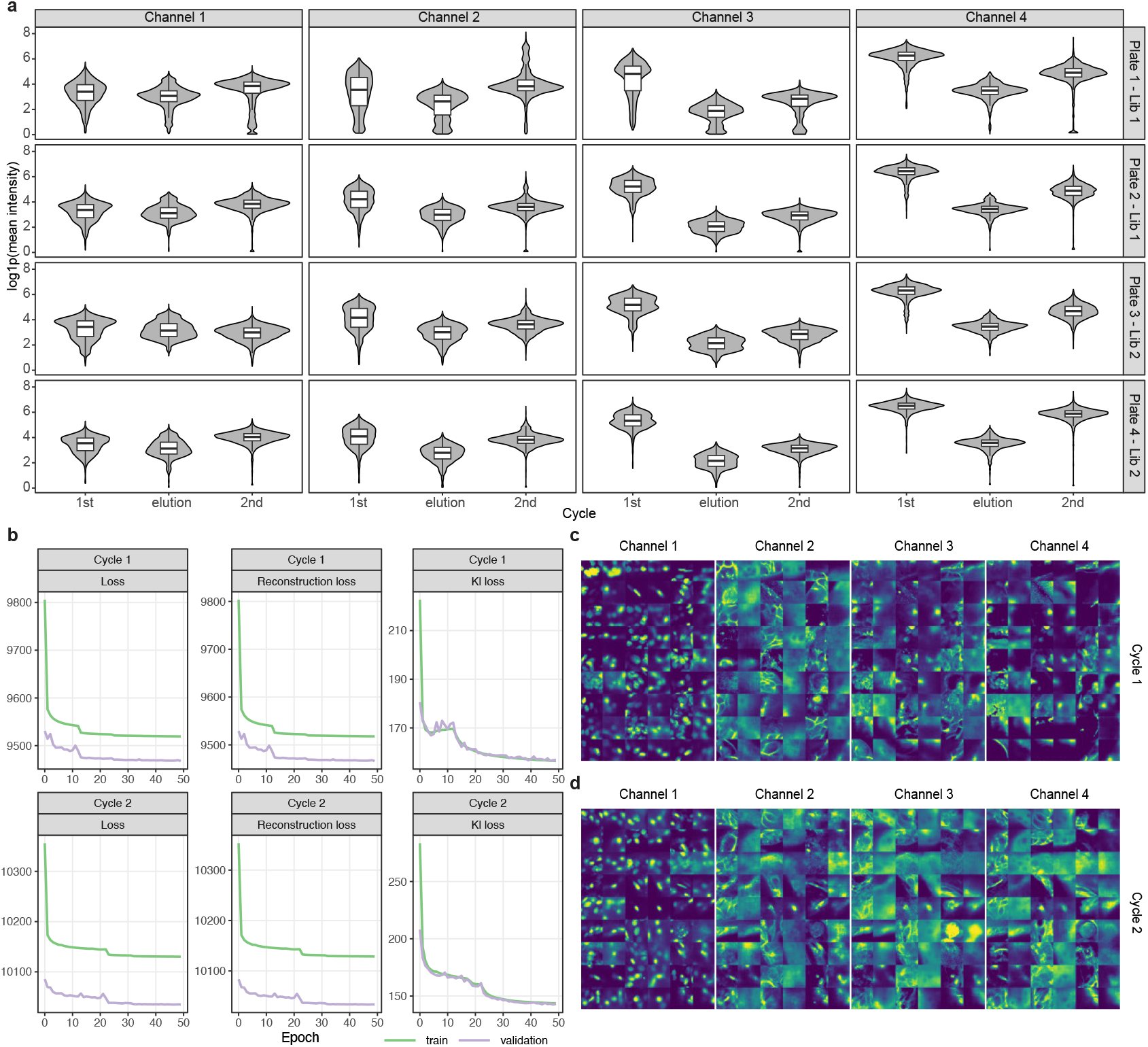
Quality control assessments for high-throughput screening dataset. a) Violin plots with overlaid boxplots showing signal intensity distributions (log mean intensity) across four fluorescence channels for each experimental phase (1st cycle, elution and 2nd cycle) in four different imaged plates. b) Training and validation loss curves for the deep learning autoencoder model used in Phenocoder. Left panels show total loss, middle panels show reconstruction loss, and right panels show KL divergence loss for both cycle 1 (top row) and cycle 2 (bottom row). c-d) Representative examples of reconstructed image patches (128×128 pixels) by the trained autoencoder model for cycle 1 (c) and cycle 2 (d) across four fluorescence channels.

**Extended Data Figure 9.**
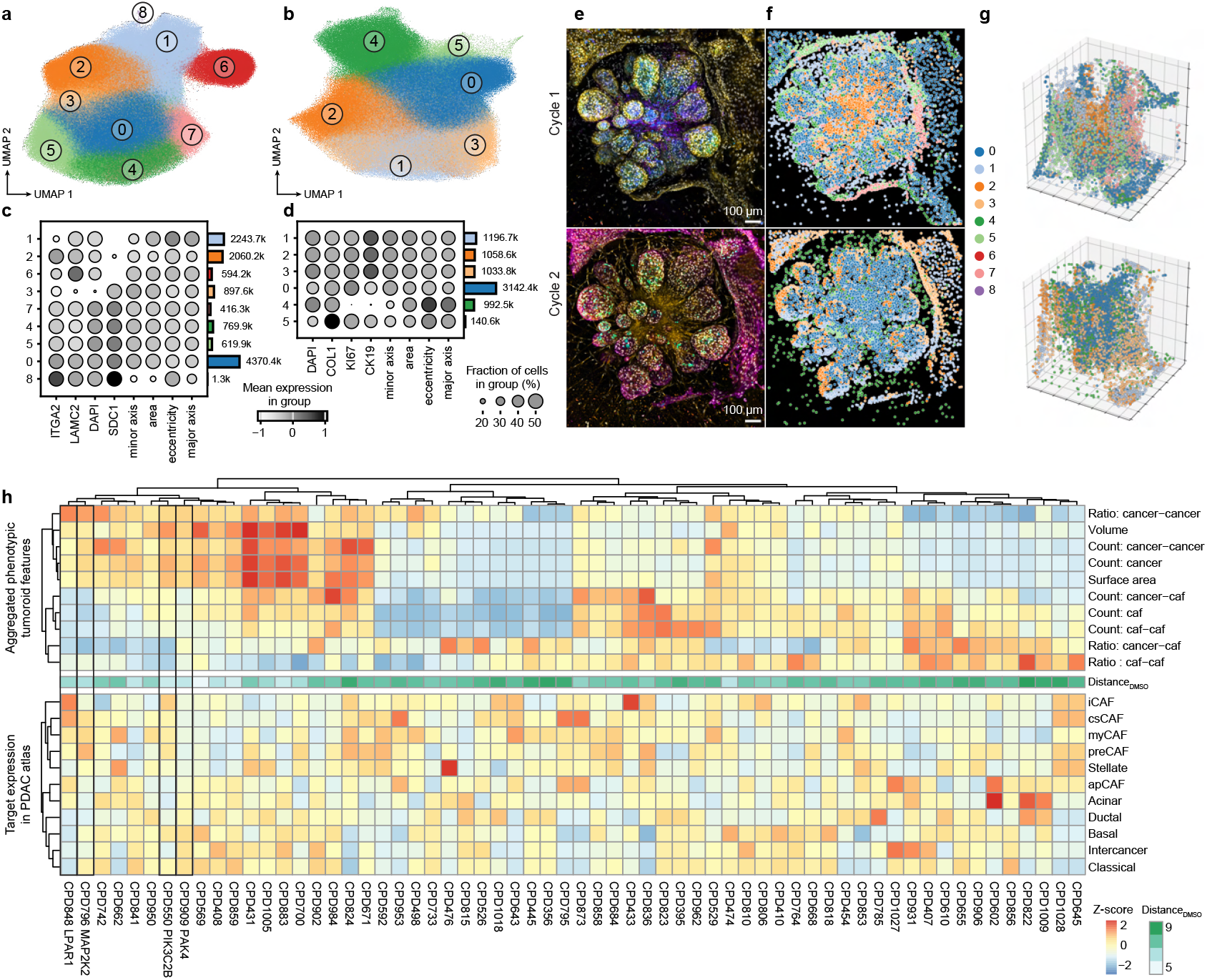
Extended analysis for systematic perturbation of cancer-CAF interactions. a-b) UMAP embeddings of single-cell phenotypic features from Phenocoder analysis of first (a) and second (b) imaging cycles with identified cell niches/environments, revealing distinct cellular microenvironments within the tumoroid cultures. c-d) Dot plots showing marker expression patterns across identified cell niches for first (c) and second (d) imaging cycles. Circle size indicates the fraction of cells in each niche expressing the marker, while color intensity represents mean expression level. Key markers include ITGA2, LAMC2, DAPI, SDC1 in the first cycle and COL1, KI67, CK19 in the second cycle as well as area, major/minor axis, eccentricity measures. e-f) Representative immunofluorescence images of a tumoroid (DMSO) (e) alongside its segmented phenotypic map (f) for both imaging cycles. g) 3D scatter plot visualization of the Phenocoder clusters, with cycle 1 shown in the top panel and cycle 2 in the bottom panel. Cell niches are color-coded to match panels a-b), providing spatial relationships between identified microenvironments in both imaging cycles. h) Top heatmap displays the effects of compound treatments on key phenotypic features (Z-score) extracted from image analysis. Features include distance from DMSO control, proliferative cluster metrics, cancer-cancer interactions, volume measurements, area quantifications, and cancer-CAF interface metrics. Bottom heatmap shows Z-scores of weighted compound target expression in cancer/CAF subtypes identified in the PDAC scRNA-seq atlas.

**Extended Data Figure 10.**
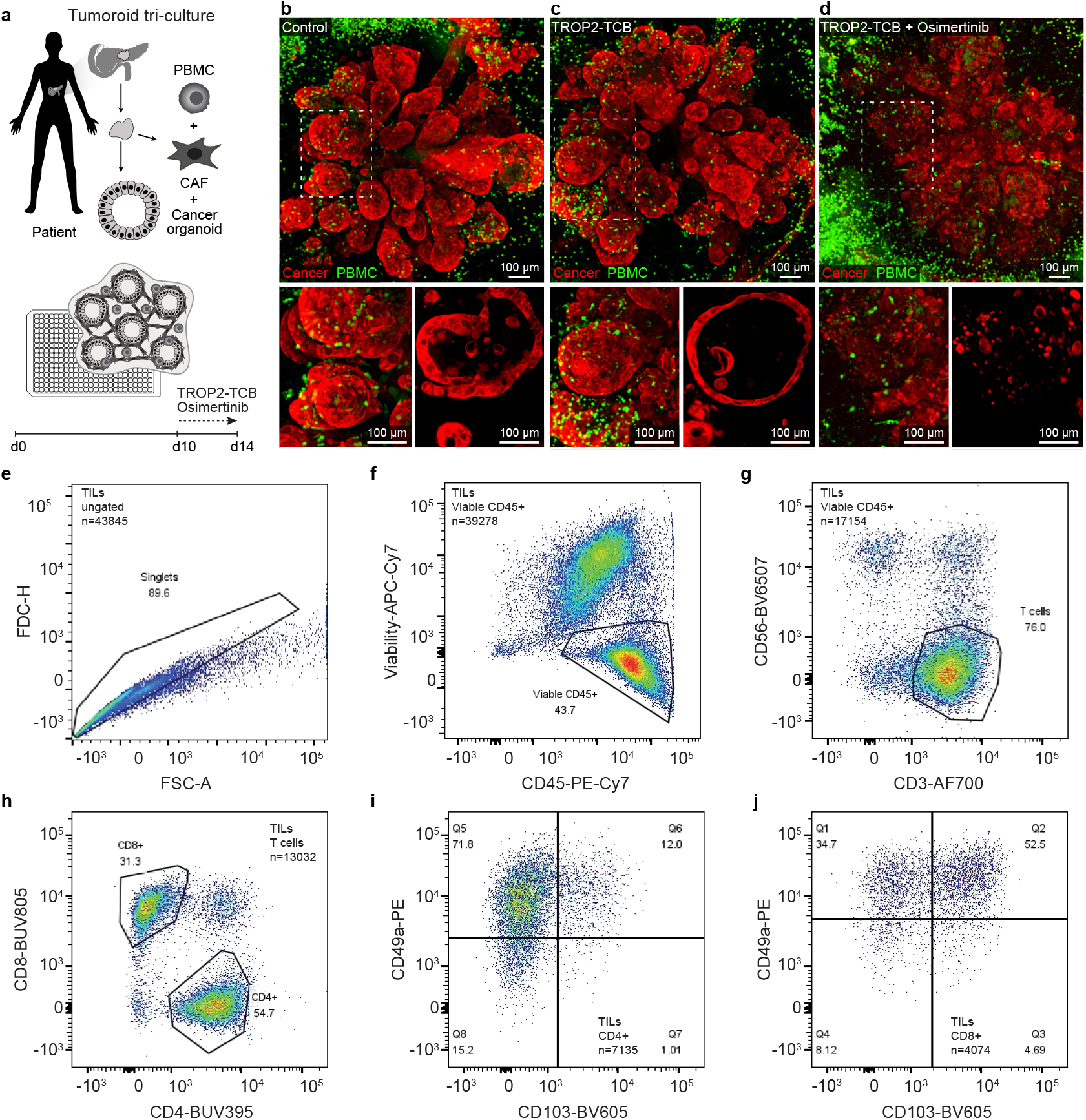
Incorporation of immune cells into cancer-CAF tumoroids. (a) Schematic overview of the high-content imaging-based phenotypic screening platform for patient-derived cancer-CAF tri-culture PDAC tumoroids including activated T cells from peripheral blood mononuclear cells (PBMCs). (b-d) 3D maximum projection (top) and z-slice (bottom) confocal images of PDAC tumoroids co-cultured with PBMC-derived T cells under the indicated treatment conditions. Cancer cells are shown in red, T cells in green. (b) Osimertinib alone, (c) Osimertinib + control TCB, and (d) Osimertinib + TROP2-TCB. e-j) Flow cytometry analysis of tumor-infiltrating lymphocytes (TILs) freshly isolated from patient-derived PDAC tissue. (e) Singlet selection, (f) viable CD45^+^ immune cells, (g) CD3^+^ T cells, (h) CD4^+^ and CD8^+^ subsets, and (i,j) further subdivision of T cell populations based on CD49a and CD103 expression.

### A. Extended Data Tables

All supplementary tables will be accessible online upon publication.

Table S1: Patient samples used in this study.

Table S2: Used antibodies.

Table S3: Compounds used in pilot screen.

Table S4: Compound library used in high throughput screening.

## References

Albanese, A., Swaney, J. M., Yun, D. H., Evans, N. B., Antonucci, J. M., Velasco, S., Sohn, C. H., Arlotta, P., Gehrke, L., and Chung, K. Multiscale 3D phenotyping of human cerebral organoids. 10:21487.

Barbazan, J., Pérez-González, C., Gómez-González, M., Dedenon, M., Richon, S., Latorre, E., Serra, M., Mariani, P., Descroix, S., Sens, P., Trepat, X., and Vignjevic, D. M. Cancer-associated fibroblasts actively compress cancer cells and modulate mechanotransduction. 14:6966.

Bardia, A., Hurvitz, S. A., Tolaney, S. M., Loirat, D., Punie, K., Oliveira, M., Brufsky, A., Sardesai, S. D., Kalinsky, K., Zelnak, A. B., Weaver, R., Traina, T., Dalenc, F., Aftimos, P., Lynce, F., Diab, S., Cortés, J., O’Shaughnessy, J., Diéras, V., Ferrario, C., Schmid, P., Carey, L. A., Gianni, L., Piccart, M. J., Loibl, S., Goldenberg, D. M., Hong, Q., Olivo, M. S., Itri, L. M., Rugo, H. S., and ASCENT Clinical Trial Investigators. Sacituzumab govitecan in metastatic triple-negative breast cancer. 384:1529–1541.

Barkovskaya, A., Kim, K., Shankar, A., Meca-Laguna, G., Rae, M., Le Saux, C. J., and Sharma, A. Inhibition of PIKfyve kinase induces senescent cell death by suppressing lysosomal exocytosis and leads to improved outcomes in a mouse model of idiopathic pulmonary fibrosis.

Beghin, A., Grenci, G., Sahni, G., Guo, S., Rajendiran, H., Delaire, T., Mohamad Raffi, S. B., Blanc, D., de Mets, R., Ong, H. T., Galindo, X., Monet, A., Acharya, V., Racine, V., Levet, F., Galland, R., Sibarita, J.-B., and Viasnoff, V. Automated high-speed 3D imaging of organoid cultures with multi-scale phenotypic quantification. 19:881–892.

Birk, S., Bonafonte-Pardàs, I., Feriz, A. M., Boxall, A., Agirre, E., Memi, F., Maguza, A., Yadav, A., Armingol, E., Fan, R., Castelo-Branco, G., Theis, F. J., Bayraktar, O. A., Talavera-López, C., and Lotfollahi, M. Quantitative characterization of cell niches in spatially resolved omics data. pages 1–13.

Boj, S. F., Hwang, C.-I., Baker, L. A., Chio, I. I. C., Engle, D. D., Corbo, V., Jager, M., Ponz-Sarvise, M., Tiriac, H., Spector, M. S., Gracanin, A., Oni, T., Yu, K. H., van Boxtel, R., Huch, M., Rivera, K. D., Wilson, J. P., Feigin, M. E., Öhlund, D., Handly-Santana, A., Ardito-Abraham, C. M., Ludwig, M., Elyada, E., Alagesan, B., Biffi, G., Yordanov, G. N., Delcuze, B., Creighton, B., Wright, K., Park, Y., Morsink, F. H. M., Molenaar, I. Q., Borel Rinkes, I. H., Cuppen, E., Hao, Y., Jin, Y., Nijman, I. J., Iacobuzio-Donahue, C., Leach, S. D., Pappin, D. J., Hammell, M., Klimstra, D. S., Basturk, O., Hruban, R. H. Okuda & Harmel et al. | Platform for high-throughput phenospace learning of 3D multilineage organoid systems | 11

Offerhaus, G. J., Vries, R. G. J., Clevers, H., and Tuveson, D. A. Organoid models of human and mouse ductal pancreatic cancer. 160:324–338.

Bredikhin, D., Kats, I., and Stegle, O. MUON: multimodal omics analysis framework. 23:42.

Caunt, C. J., Sale, M. J., Smith, P. D., and Cook, S. J. MEK1 and MEK2 inhibitors and cancer therapy: the long and winding road. 15:577–592.

Chen, M.-M., Gao, Q., Ning, H., Chen, K., Gao, Y., Yu, M., Liu, C.-Q., Zhou, W., Pan, J., Wei, L., Dou, W., Zhang, D., Zhu, L., Zhang, Q., Chen, R., and Zhang, Z. Integrated single-cell and spatial transcriptomics uncover distinct cellular subtypes involved in neural invasion in pancreatic cancer. 43:1656–1676.e10.

Cheng, C., Hu, J., Mannan, R., He, T., Bhattacharyya, R., Magnuson, B., Wisniewski, J. P., Peters, S., Karim, S. A., MacLean, D. J., Karabürk, H., Zhang, L., Rossiter, N. J., Zheng, Y., Xiao, L., Li, C., Awad, D., Mahapatra, S., Bao, Y., Zhang, Y., Cao, X., Wang, Z., Mehra, R., Morlacchi, P., Sahai, V., Pasca di Magliano, M., Shah, Y. M., Weisman, L. S., Morton, J. P., Ding, K., Qiao, Y., Lyssiotis, C. A., and Chinnaiyan, A. M. Targeting PIKfyve-driven lipid metabolism in pancreatic cancer.

Cheng, P. S. W., Zaccaria, M., and Biffi, G. Functional heterogeneity of fibroblasts in primary tumors and metastases. 11:135–153.

Cinato, M., Guitou, L., Saidi, A., Timotin, A., Sperazza, E., Duparc, T., Zolov, S. N., Giridharan, S. S. P., Weisman, L. S., Martinez, L. O., Roncalli, J., Kunduzova, O., Tronchere, H., and Boal, F. Apilimod alters TGFβ signaling pathway and prevents cardiac fibrotic remodeling. 11:6491–6506.

Corte, T. J., Behr, J., Cottin, V., Glassberg, M. K., Kreuter, M., Martinez, F. J., Ogura, T., Suda, T., Wijsenbeek, M., Berkowitz, E., Elpers, B., Kim, S., Watanabe, H., Fischer, A., and Maher, T. M. Efficacy and safety of admilparant, an LPA1 antagonist, in pulmonary fibrosis: A phase 2 randomized clinical trial. 211:230–238.

Cutrona, M. B., Wu, J., Yang, K., Peng, J., and Chen, T. Pancreatic cancer organoid-screening captures personalized sensitivity and chemoresistance suppression upon cytochrome P450 3A5-targeted inhibition. 27:110289.

Dong, W., Moses, C., and Li, K. Efficient k-nearest neighbor graph construction for generic similarity measures. In Proceedings of the 20th international conference on World wide web. ACM.

Dreyer, S. B., Beer, P., Hingorani, S. R., and Biankin, A. V. Improving outcomes of patients with pancreatic cancer. 22:439–456.

Driehuis, E., Gracanin, A., Vries, R. G. J., Clevers, H., and Boj, S. F. Establishment of pancreatic organoids from normal tissue and tumors. 1:100192.

Driehuis, E., van Hoeck, A., Moore, K., Kolders, S., Francies, H. E., Gulersonmez, M. C., Stigter, E. C. A., Burgering, B., Geurts, V., Gracanin, A., Bounova, G., Morsink, F. H., Vries, R., Boj, S., van Es, J., Offerhaus, G. J. A., Kranenburg, O., Garnett, M. J., Wessels, L., Cuppen, E., Brosens, L. A. A., and Clevers, H. Pancreatic cancer organoids recapitulate disease and allow personalized drug screening. 116:26580–26590.

Fleck, J. S., Camp, J. G., and Treutlein, B. What is a cell type? 381:733–734.

Gao, Y., Li, J., Cheng, W., Diao, T., Liu, H., Bo, Y., Liu, C., Zhou, W., Chen, M., Zhang, Y., Liu, Z., Han, W., Chen, R., Peng, J., Zhu, L., Hou, W., and Zhang, Z. Cross-tissue human fibroblast atlas reveals myofibroblast subtypes with distinct roles in immune modulation. 42:1764–1783.e10.

Grünwald, B. T., Devisme, A., Andrieux, G., Vyas, F., Aliar, K., McCloskey, C. W., Macklin, A., Jang, G. H., Denroche, R., Romero, J. M., Bavi, P., Bronsert, P., Notta, F., O’Kane, G., Wilson, J., Knox, J., Tamblyn, L., Udaskin, M., Radulovich, N., Fischer, S. E., Boerries, M., Gallinger, S., Kislinger, T., and Khokha, R. Spatially confined sub-tumor microenvironments in pancreatic cancer. 184:5577–5592.e18.

Gut, G., Herrmann, M. D., and Pelkmans, L. Multiplexed protein maps link subcellular organization to cellular states. 361(6401).

Haghverdi, L., Buettner, F., and Theis, F. J. Diffusion maps for high-dimensional single-cell analysis of differentiation data. 31(18):2989–2998.

Halbrook, C. J., Lyssiotis, C. A., Pasca di Magliano, M., and Maitra, A. Pancreatic cancer: Advances and challenges. 186:1729–1754.

Herrera, M., Pretelli, G., Desai, J., Garralda, E., Siu, L. L., Steiner, T. M., and Au, L. Bispecific antibodies: advancing precision oncology. 10:893–919.

Hirt, C. K., Booij, T. H., Grob, L., Simmler, P., Toussaint, N. C., Keller, D., Taube, D., Ludwig, V., Goryachkin, A., Pauli, C., Lenggenhager, D., Stekhoven, D. J., Stirnimann, C. U., Endhardt, K., Ringnalda, F., Villiger, L., Siebenhüner, A., Karkampouna, S., De Menna, M., Beshay, J., Klett, H., Kruithof-de Julio, M., Schüler, J., and Schwank, G. Drug screening and genome editing in human pancreatic cancer organoids identifies drug-gene interactions and candidates for off-label treatment. 2:100095.

Hwang, W. L., Jagadeesh, K. A., Guo, J. A., Hoffman, H. I., Yadollahpour, P., Reeves, J. W., Mohan, R., Drokhlyansky, E., Van Wittenberghe, N., Ashenberg, O., Farhi, S. L., Schapiro, D., Divakar, P., Miller, E., Zollinger, D. R., Eng, G., Schenkel, J. M., Su, J., Shiau, C., Yu, P., Freed-Pastor, W. A., Abbondanza, D., Mehta, A., Gould, J., Lambden, C., Porter, C. B. M., Tsankov, A., Dionne, D., Waldman, J., Cuoco, M. S., Nguyen, L., Delorey, T., Phillips, D., Barth, J. L., Kem, M., Rodrigues, C., Ciprani, D., Roldan, J., Zelga, P., Jorgji, V., Chen, J. H., Ely, Z., Zhao, D., Fuhrman, K., Fropf, R., Beechem, J. M., Loeffler, J. S., Ryan, D. P., Weekes, C. D., Ferrone, C. R., Qadan, M., Aryee, M. J., Jain, R. K., Neuberg, D. S., Wo, J. Y., Hong, T. S., Xavier, R., Aguirre, A. J., Rozenblatt-Rosen, O., Mino-Kenudson, M., Castillo, C. F.-D., Liss, A. S., Ting, D. T., Jacks, T., and Regev, A. Single-nucleus and spatial transcriptome profiling of pancreatic cancer identifies multicellular dynamics associated with neoadjuvant treatment. 54:1178–1191.

Juin, A., Spence, H. J., Martin, K. J., McGhee, E., Neilson, M., Cutiongco, M. F. A., Gadegaard, N., Mackay, G., Fort, L., Lilla, S., Kalna, G., Thomason, P., Koh, Y. W. H., Norman, J. C., Insall, R. H., and Machesky, L. M. N-WASP control of LPAR1 trafficking establishes response to self-generated LPA gradients to promote pancreatic cancer cell metastasis. 51:431–445.e7.

Kim, E., Choi, S., Kang, B., Kong, J., Kim, Y., Yoon, W. H., Lee, H.-R., Kim, S., Kim, H.M., Lee, H., Yang, C., Lee, Y. J., Kang, M., Roh, T.-Y., Jung, S., Kim, S., Ku, J. H., and Shin, K. Creation of bladder assembloids mimicking tissue regeneration and cancer. 588: 664–669.

Kim, J., Rustam, S., Mosquera, J. M., Randell, S. H., Shaykhiev, R., Rendeiro, A. F., and Elemento, O. Unsupervised discovery of tissue architecture in multiplexed imaging. 19:

Okuda & Harmel et al. | Platform for high-throughput phenospace learning of 3D multilineage organoid systems 1653–1661.

Kingma, D. P. and Welling, M. Auto-encoding variational bayes.

Klein, C., Brinkmann, U., Reichert, J. M., and Kontermann, R. E. The present and future of bispecific antibodies for cancer therapy. 23:301–319.

Langer, E. M., Allen-Petersen, B. L., King, S. M., Kendsersky, N. D., Turnidge, M. A., Kuziel, G. M., Riggers, R., Samatham, R., Amery, T. S., Jacques, S. L., Sheppard, B. C., Korkola, J. E., Muschler, J. L., Thibault, G., Chang, Y. H., Gray, J. W., Presnell, S. C., Nguyen, D. G., and Sears, R. C. Modeling tumor phenotypes in vitro with three-dimensional bioprinting. 26:608–623.e6.

Li, Q., Chen, Q., Wang, W., Xie, R., Li, Z., and Chen, D. KGF secreted from HSCs activates PAK4/BMI1, promotes HCC stemness through PI3K/AKT pathway. 77:e2929.

Li, Y., Tang, S., Wang, H., Zhu, H., Lu, Y., Zhang, Y., Guo, S., He, J., Li, Y., Zhang, Y., Shi, X., Miao, Y., Zhong, C., Zhu, Y., Ju, Y., Liu, Y., Sun, M., Wang, Y., Chen, L., Zhou, H., Jin, G., and Gao, D. A pancreatic cancer organoid biobank links multi-omics signatures to therapeutic response and clinical evaluation of statin combination therapy. 32:1369– 1389.e14.

Liu, H., Noguera-Ortega, E., Dong, X., Lee, W. D., Chang, J., Aydin, S. A., Li, Y., Shin, Y., Shi, X., Liousia, M., Martinez, M. C., Brotman, J. J., Kim, S., Chen, Z., Wang, A., Ou, Z., Paek, J., Park, J. Y., Liu, A., Hu, H., Xiao, Z., Racca, D. M., Kim, S.-J., Worthen, G. S., Guo, W., Puré, E., Kang, T., Rabinowitz, J. D., Wherry, E. J., Moon, E. K., Albelda, S. M., and Huh, D. D. A tumor-on-a-chip for in vitro study of CAR-T cell immunotherapy in solid tumors.

Liu, Y., Sinjab, A., Min, J., Han, G., Paradiso, F., Zhang, Y., Wang, R., Pei, G., Dai, Y., Liu, Y., Cho, K. S., Dai, E., Basi, A., Burks, J. K., Rajapakshe, K. I., Chu, Y., Jiang, J., Zhang, D., Yan, X., Guerrero, P. A., Serrano, A., Li, M., Hwang, T. H., Futreal, A., Ajani, J. A., Solis Soto, L. M., Jazaeri, A. A., Kadara, H., Maitra, A., and Wang, L. Conserved spatial subtypes and cellular neighborhoods of cancer-associated fibroblasts revealed by single-cell spatial multi-omics.

Lukonin, I., Serra, D., Challet Meylan, L., Volkmann, K., Baaten, J., Zhao, R., Meeusen, S., Colman, K., Maurer, F., Stadler, M. B., Jenkins, J., and Liberali, P. Phenotypic landscape of intestinal organoid regeneration. 586:275–280.

Metzger, J. J., Pereda, C., Adhikari, A., Haremaki, T., Galgoczi, S., Siggia, E. D., Brivanlou, A. H., and Etoc, F. Deep-learning analysis of micropattern-based organoids enables high-throughput drug screening of huntington’s disease models. 2:100297.

Monteran, L., Zait, Y., and Erez, N. It’s all about the base: stromal cells are central orchestrators of metastasis. 10:208–229.

Nguyen, T. K., Tata, P., Brooks, S., Jena, N., Morse, S. J., Liu, Z.-Y., Lai, H. Y., Rezk, S., Van Etten, R. A., and Fleischman, A. G. The MEK/ERK inhibitor trametinib reduces fibrosis in a transduction-transplantation model of mutated calreticulin. 128:635–635.

Ong, H. T., Karatas, E., Poquillon, T., Grenci, G., Furlan, A., Dilasser, F., Mohamad Raffi, S. B., Blanc, D., Drimaracci, E., Mikec, D., Galisot, G., Johnson, B. A., Liu, A. Z., Thiel, C., Ullrich, O., Racine, V., and Beghin, A. Digitalized organoids: integrated pipeline for high-speed 3D analysis of organoid structures using multilevel segmentation and cellular topology. pages 1–12.

Palikuqi, B., Nguyen, D.-H. T., Li, G., Schreiner, R., Pellegata, A. F., Liu, Y., Redmond, D., Geng, F., Lin, Y., Gómez-Salinero, J. M., Yokoyama, M., Zumbo, P., Zhang, T., Kunar, B., Witherspoon, M., Han, T., Tedeschi, A. M., Scottoni, F., Lipkin, S. M., Dow, L., Elemento, O., Xiang, J. Z., Shido, K., Spence, J. R., Zhou, Q. J., Schwartz, R. E., De Coppi, P., Rabbany, S. Y., and Rafii, S. Adaptable haemodynamic endothelial cells for organogenesis and tumorigenesis. 585:426–432.

Palla, G., Spitzer, H., Klein, M., Fischer, D., Schaar, A. C., Kuemmerle, L. B., Rybakov, S., Ibarra, I. L., Holmberg, O., Virshup, I., Lotfollahi, M., Richter, S., and Theis, F. J. Squidpy: a scalable framework for spatial omics analysis. 19:171–178.

Pei, G., Min, J., Rajapakshe, K. I., Branchi, V., Liu, Y., Selvanesan, B. C., Thege, F., Sadeghian, D., Zhang, D., Cho, K. S., Chu, Y., Dai, E., Han, G., Li, M., Yee, C., Takahashi, K., Garg, B., Tiriac, H., Bernard, V., Semaan, A., Grem, J. L., Caffrey, T. C., Burks, J. K., Lowy, A. M., Aguirre, A. J., Grandgenett, P. M., Hollingsworth, M. A., Guerrero, P. A., Wang, L., and Maitra, A. Spatial mapping of transcriptomic plasticity in metastatic pancreatic cancer.

Peng, J., Sun, B.-F., Chen, C.-Y., Zhou, J.-Y., Chen, Y.-S., Chen, H., Liu, L., Huang, D., Jiang, J., Cui, G.-S., Yang, Y., Wang, W., Guo, D., Dai, M., Guo, J., Zhang, T., Liao, Q., Liu, Y., Zhao, Y.-L., Han, D.-L., Zhao, Y., Yang, Y.-G., and Wu, W. Single-cell RNA-seq highlights intra-tumoral heterogeneity and malignant progression in pancreatic ductal adenocarcinoma. 29:725–738.

Peng, T., Thorn, K., Schroeder, T., Wang, L., Theis, F. J., Marr, C., and Navab, N. A BaSiC tool for background and shading correction of optical microscopy images. 8.

Polański, K., Young, M. D., Miao, Z., Meyer, K. B., Teichmann, S. A., and Park, J.-E. BBKNN: fast batch alignment of single cell transcriptomes. 36:964–965.

Provenzano, P. P., Cuevas, C., Chang, A. E., Goel, V. K., Von Hoff, D. D., and Hingorani, S. R. Enzymatic targeting of the stroma ablates physical barriers to treatment of pancreatic ductal adenocarcinoma. 21:418–429.

Punovuori, K., Bertillot, F., Miroshnikova, Y. A., Binner, M. I., Myllymäki, S.-M., Follain, G., Kruse, K., Routila, J., Huusko, T., Pellinen, T., Hagström, J., Kedei, N., Ventelä, S., Mäkitie, A., Ivaska, J., and Wickström, S. A. Multiparameter imaging reveals clinically relevant cancer cell-stroma interaction dynamics in head and neck cancer. 187:7267–7284.e20.

Raffo-Romero, A., Ziane-Chaouche, L., Salomé-Desnoulez, S., Hajjaji, N., Fournier, I., Salzet, M., and Duhamel, M. A co-culture system of macrophages with breast cancer tumoroids to study cell interactions and therapeutic responses. 4:100792.

Raghavan, S., Winter, P. S., Navia, A. W., Williams, H. L., DenAdel, A., Lowder, K. E., Galvez-Reyes, J., Kalekar, R. L., Mulugeta, N., Kapner, K. S., Raghavan, M. S., Borah, A. A., Liu, N., Väyrynen, S. A., Costa, A. D., Ng, R. W. S., Wang, J., Hill, E. K., Ragon, D. Y., Brais, L. K., Jaeger, A. M., Spurr, L. F., Li, Y. Y., Cherniack, A. D., Booker, M. A., Cohen, E. F., Tolstorukov, M. Y., Wakiro, I., Rotem, A., Johnson, B. E., McFarland, J. M., Sicinska, E. T., Jacks, T. E., Sullivan, R. J., Shapiro, G. I., Clancy, T. E., Perez, K., Rubinson, D. A., Ng, K., Cleary, J. M., Crawford, L., Manalis, S. R., Nowak, J. A., Wolpin, B. M., Hahn, W. C., Aguirre, A. J., and Shalek, A. K. Microenvironment drives cell state, plasticity, and drug response in pancreatic cancer. 184:6119–6137.e26.

Ramos Zapatero, M., Tong, A., Opzoomer, J. W., O’Sullivan, R., Cardoso Rodriguez, F., Sufi, J., Vlckova, P., Nattress, C., Qin, X., Claus, J., Hochhauser, D., Krishnaswamy, S., and Tape, C. J. Trellis tree-based analysis reveals stromal regulation of patient-derived organoid drug responses. 186:5606–5619.e24.

Recaldin, T., Steinacher, L., Gjeta, B., Harter, M. F., Adam, L., Kromer, K., Mendes, M. P., Bellavista, M., Nikolaev, M., Lazzaroni, G., Krese, R., Kilik, U., Popovic, D., Stoll, B., Gerard, R., Bscheider, M., Bickle, M., Cabon, L., Camp, J. G., and Gjorevski, N. Human organoids with an autologous tissue-resident immune compartment. 633:165–173.

Rocklin, M. Dask: Parallel computation with blocked algorithms and task scheduling. In Proceedings of the Python in Science Conference, pages 126–132. SciPy.

Rood, J. E., Hupalowska, A., and Regev, A. Toward a foundation model of causal cell and tissue biology with a perturbation cell and tissue atlas. 187:4520–4545.

Seino, T., Kawasaki, S., Shimokawa, M., Tamagawa, H., Toshimitsu, K., Fujii, M., Ohta, Y., Matano, M., Nanki, K., Kawasaki, K., Takahashi, S., Sugimoto, S., Iwasaki, E., Takagi, J., Itoi, T., Kitago, M., Kitagawa, Y., Kanai, T., and Sato, T. Human pancreatic tumor organoids reveal loss of stem cell niche factor dependence during disease progression. 22:454–467.e6.

Sharpe, B. P., Nazlamova, L. A., Tse, C., Johnston, D. A., Thomas, J., Blyth, R., Pickering, O. J., Grace, B., Harrington, J., Rajak, R., Rose-Zerilli, M., Walters, Z. S., and Underwood, T. J. Patient-derived tumor organoid and fibroblast assembloid models for interrogation of the tumor microenvironment in esophageal adenocarcinoma. 4:100909.

Siegel, R. L., Kratzer, T. B., Giaquinto, A. N., Sung, H., and Jemal, A. Cancer statistics, 2025. 75:10–45.

Stringer, C. and Pachitariu, M. Cellpose3: one-click image restoration for improved cellular segmentation. 22:592–599.

Szklarczyk, D., Kirsch, R., Koutrouli, M., Nastou, K., Mehryary, F., Hachilif, R., Gable, A. L., Fang, T., Doncheva, N. T., Pyysalo, S., Bork, P., Jensen, L. J., and von Mering, C. The STRING database in 2023: protein-protein association networks and functional enrichment analyses for any sequenced genome of interest. 51:D638–D646.

Verstegen, M. M. A., Coppes, R. P., Beghin, A., De Coppi, P., Gerli, M. F. M., de Graeff, N., Pan, Q., Saito, Y., Shi, S., Zadpoor, A. A., and van der Laan, L. J. W. Clinical applications of human organoids. 31:409–421.

Volat, F., Medhi, R., Maggs, L. Z., Deken, M. A., Price, A., Andrews, L., Clark, J., Taylor, D., Carruthers, A., Taylor-Smith, E., Pacheco, N., Rudge, S. A., Fraser, A., Lopez-Clavijo, A. F., Sousa, B. C., Johnson, Z., Di Conza, G., van der Veen, L., Shah, P., Sandig, H., Sharpe, H. J., and Farrow, S. Pancreatic CAF-derived autotaxin drives autocrine CTGF expression to modulate protumorigenic signaling. 24:230–241.

Volkmann, E. R., Denton, C. P., Kolb, M., Wijsenbeek-Lourens, M. S., Emson, C., Hudson, K., Amatucci, A. J., Distler, O., Allanore, Y., and Khanna, D. Lysophosphatidic acid receptor 1 inhibition: a potential treatment target for pulmonary fibrosis. 33:240015.

Wahle, P., Brancati, G., Harmel, C., He, Z., Gut, G., Del Castillo, J. S., Xavier da Silveira Dos Santos, A., Yu, Q., Noser, P., Fleck, J. S., Gjeta, B., Pavlinić, D., Picelli, S., Hess, M., Schmidt, G. W., Lummen, T. T. A., Hou, Y., Galliker, P., Goldblum, D., Balogh, M., Cowan, C. S., Scholl, H. P. N., Roska, B., Renner, M., Pelkmans, L., Treutlein, B., and Camp, J. G. Multimodal spatiotemporal phenotyping of human retinal organoid development.

Wolf, F. A., Angerer, P., and Theis, F. J. SCANPY: large-scale single-cell gene expression data analysis. 19:15.

Xu, Q., Okuda, R., Gjeta, B., Harmel, C., Signer, M., Lucioli, M., Santel, M., Seimiya, M., Esposito, C., Guja-Jarosz, K., Maynard, A., Morinaga, S., Miyagi, Y., Yamaguchi, T., Ueno, Y., Piscuoglio, S., Müller, D. J., Taniguichi, H., Treutlein, B., and Camp, J. G. Integrated cell atlas and tumoroids chart pancreatic cancer therapeutic targets. page 2025.09.30.679518.

Xu, S., Ma, B., Jian, Y., Yao, C., Wang, Z., Fan, Y., Ma, J., Chen, Y., Feng, X., An, J., Chen, J., Wang, K., Xie, H., Gao, Y., and Li, L. Development of a PAK4-targeting PROTAC for renal carcinoma therapy: concurrent inhibition of cancer cell proliferation and enhancement of immune cell response. 104:105162.

Zapatero, M. R., Tong, A., Opzoomer, J. W., O’Sullivan, R., Rodriguez, F. C., Sufi, J., Vlckova, P., Nattress, C., Qin, X., Claus, J., Hochhauser, D., Krishnaswamy, S., and Tape, C. J. Trellis tree-based analysis reveals stromal regulation of patient-derived organoid drug responses. 187:7335–7349.

